# The Carotid Body Detects Circulating Tumor Necrosis Factor-Alpha to Activate a Sympathetic Anti-Inflammatory Reflex

**DOI:** 10.1101/2021.10.20.463417

**Authors:** Pedro L. Katayama, Isabela P. Leirão, Alexandre Kanashiro, João Paulo M. Luiz, Fernando Q. Cunha, Luiz C. C. Navegantes, Jose V. Menani, Daniel B. Zoccal, Débora S. A. Colombari, Eduardo Colombari

## Abstract

Recent evidence has suggested that the carotid bodies might act as immunological sensors, detecting pro-inflammatory mediators and signalling to the central nervous system, which, in turn, orchestrates autonomic responses. Here, we demonstrated that the TNF-α receptor type I is expressed in the carotid bodies of rats. The systemic administration of TNF-α increased carotid body afferent discharge and activated glutamatergic neurons in the nucleus tractus solitarius (NTS) that project to the rostral ventrolateral medulla (RVLM), where the majority of pre-sympathetic neurons reside. The activation of these neurons was accompanied by generalized activation of the sympathetic nervous system. Carotid body ablation blunted the TNF-α-induced activation of RVLM-projecting NTS neurons and the increase in splanchnic sympathetic nerve activity. Finally, plasma and spleen levels of cytokines after TNF-α administration were higher in rats subjected to either carotid body ablation or splanchnic sympathetic denervation. Collectively, our findings indicate that the carotid body detects circulating TNF-α to activate a counteracting sympathetic anti-inflammatory mechanism.

## Introduction

The existence of neuroimmune interactions and their relevance to the control of inflammation are well-established and have been extensively explored in the last 20 years (Abe et al., 2017; Araujo et al., 2019; Bassi et al., 2020; Chavan et al., 2017; Chu et al., 2020; Gabanyi et al., 2016; Kressel et al., 2020; Lankadeva et al., 2020; Martelli et al., 2014; Murray et al., 2021; Steinman, 2004; Tanaka et al., 2021) since the discovery of the “inflammatory reflex” (Borovikova et al., 2000). In general, there is a consensus that this reflex works as a negative-feedback mechanism that comprises: 1) a detection component, which identifies pathogen- or danger-associated molecular patterns, generating an inflammatory response; 2) an afferent arm, which conveys information about the systemic inflammatory status to the central nervous system; 3) integrative centers in the brain, that receive and process signals regarding the systemic inflammatory condition, orchestrating an appropriate counteracting response and; 4) an efferent arm, which are the effectors that exert immunomodulatory functions to promote resolution of infection and inflammation.

The vagus nerve is considered an important element in neuroimmune interactions (Borovikova et al., 2000; Kressel et al., 2020). Its afferent (sensory) and efferent (motor) fibers are involved in the bidirectional communication between the nervous and the immune systems, providing a reflex mechanism known as the “cholinergic anti-inflammatory pathway” (Borovikova et al., 2000). According this mechanism, vagal sensory neurons detect inflammatory mediators produced in conditions of systemic inflammation and send this information to the central nervous system (Watkins et al., 1995), which, in turn, generates a vagal efferent output that counteracts inflammation mainly through acetylcholine-induced inhibition of cytokine production (Borovikova et al., 2000). The importance of this cholinergic anti-inflammatory mechanism is beyond doubt since its dysfunction is involved in the pathophysiology of several conditions (Bassi et al., 2017; Chang et al., 2019; Kanashiro et al., 2017; Li et al., 2011; van Maanen et al., 2009). However, several studies have shown convincing evidence for the existence of other neural mechanisms that regulate inflammation. For instance, animal and human studies have demonstrated that the efferent sympathetic nervous system can modulate inflammatory conditions through catecholamine-mediated suppression of innate immune responses (Abe et al., 2017; Kox et al., 2014; Lankadeva et al., 2020; Martelli et al., 2014; Tanaka et al., 2021; van Westerloo et al., 2011). Moreover, some studies demonstrated that the sympathetic-mediated anti-inflammatory reflexes do not depend on vagal afferent signalling, suggesting the existence of other peripheral mechanisms able to detect inflammation and communicate with the central nervous system to activate downstream sympathetic anti-inflammatory pathways (Abe et al., 2017; Martelli et al., 2014).

In this regard, the carotid body, classically known as the main peripheral monitor of the O_2_ levels in the blood, has been considered a polymodal sensor due to its particular ability to detect diverse molecules present in the circulation, such as glucose, sodium chloride, hormones, and also, inflammatory mediators (Allen, 1998; da Silva et al., 2019; Jendzjowsky et al., 2018; Katayama, 2016; Kumar and Prabhakar, 2012; Thompson et al., 2016). In the context of inflammation, several pieces of evidence indicate that the carotid bodies might be involved in the intricate interplay between the immune system and the sympathetic nervous system. First, the carotid body expresses receptors for inflammatory mediators such as lysophosphatidic acid (LPA) and pro-inflammatory cytokines such as IL-1β, IL-6, and tumor necrosis factor-alpha (TNF-α) (Fernández et al., 2008; Jendzjowsky et al., 2018; Kumar and Prabhakar, 2012; Mkrtchian et al., 2012; Wang et al., 2002). Second, LPA and pro-inflammatory cytokines stimulate the carotid body and increase the carotid sinus nerve (CSN) afferent activity in isolated *in vitro* preparations (Jendzjowsky et al., 2021, 2018). Third, carotid body stimulation by its typical stimulus (hypoxia) activates central autonomic areas that control parasympathetic (Erickson and Millhorn, 1994; Zera et al., 2019) and, also, the sympathetic nervous system (Kline et al., 2010; Koshiya and Guyenet, 1996; Luise King et al., 2012) which, besides vagally-mediated mechanisms, represents an important component in the neural regulation of immunity (Abe et al., 2017; Lankadeva et al., 2020; Martelli et al., 2014). Last, carotid body denervation worsens systemic inflammation and accelerates multiple organ dysfunction and death in rats with lipopolysaccharide (LPS)-induced sepsis (Nardocci et al., 2015), suggesting that the carotid body is a protective factor during acute inflammatory conditions. Altogether, these observations led to the hypothesis that the carotid body plays a role in neuroimmune interactions, but the exact mechanisms underlying this cross-talk are largely unknown.

In this study, we focused on investigating the impact of TNF-α (a ubiquitous cytokine that triggers inflammation)(Grieve et al., 2017) on the carotid body-mediated activation of the sympathetic nervous system, as well as the relevance of this interaction in the modulation of TNF-α-induced systemic inflammation. We revealed that the carotid body expresses the TNF-α receptor type I (TNFR1) and detects increased levels of TNF-α in peripheral circulation, transmitting this information to the brain via CSN afferent inputs to commissural nucleus tractus solitarius (cNTS) glutamatergic neurons that project to rostral ventrolateral medulla (RVLM) pre-sympathetic neurons, resulting in activation of the sympathetic nervous system to counteract the TNF-α-induced inflammation. We, therefore, propose the existence of a physiological carotid body-mediated neuroimmune reflex that acutely controls inflammation. The identification of this neuroimmune reflex provides potential mechanistic insights into the pathophysiology of inflammation-mediated diseases as well as into the development of novel therapeutic strategies to treat these conditions.

## Results

### TNFR1 is expressed in the carotid body

The expression of TNF-α receptors type I (TNFR1) in the carotid body was verified using two different methods: 1) RNAscope *in situ* hybridization for labelling TNFR1 mRNA molecules combined with immunofluorescence staining for tyrosine hydroxylase (TH) to identify carotid body glomus cells (Fig. 1A – C) and; 2) double immunofluorescence staining for TNFR1 and TH (Fig. 1D – F). We found that the TNFR1 is expressed in the carotid body of rats at both mRNA and protein levels (Fig. 1).

**Figure 1.**
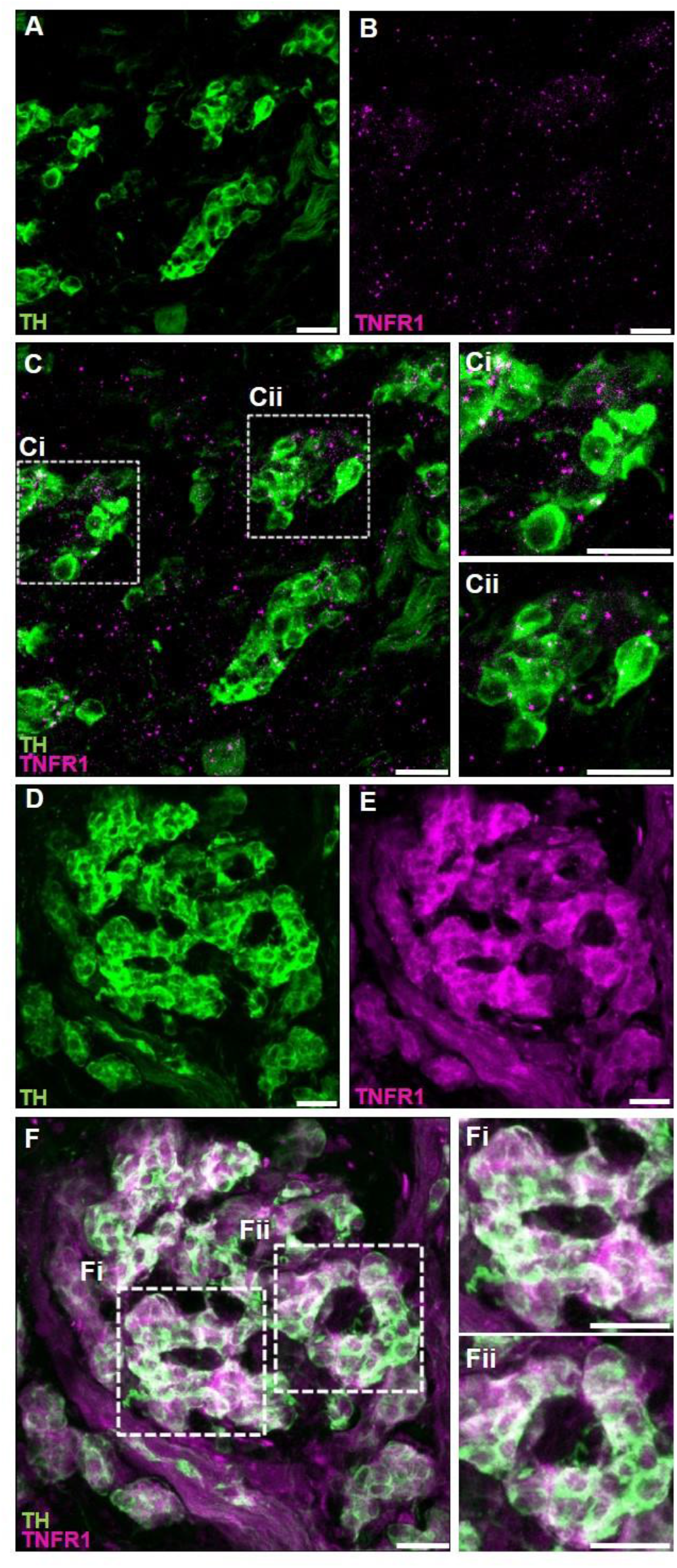
TNF-α receptors type I (TNFR1) are expressed in the carotid body of rats. **A – C.** Combined fluorescent *in situ* hybridization (TNFR1, magenta puncta) and immunofluorescence (Tyrosine hydroxylase, TH, green staining). **A.** TH positive cells (glomus cells) in the carotid body. **B.** RNAscope *in situ* hybridization showing TNFR1 mRNA expression in the carotid body. **C.** Overlay of images A and B showing the colocalization of TH and TNFR1. **Ci** and **Cii.** Zoom into selected regions of image C. **D – F.** Double immunofluorescence staining (TNFR1, magenta staining; and TH, green staining). **D.** TH positive cells (glomus cells) in the carotid body. **E.** TNFR1 expression in the carotid body. **F.** Overlay of images D and E showing the colocalization of TH and TNFR1. **Fi** and **Fii.** Zoom into selected regions of image F. Scale bars: 20 µm.

### Circulating TNF-α increases carotid sinus nerve afferent activity

Next, we investigated if elevated TNF-α levels in the blood could activate its receptors in the carotid body and increase CSN activity in vivo. We found that exogenous TNF-α administration increased CSN activity by 34 ± 5% at 30 minutes after administration compared to baseline (Figure 2B). This TNF-α-induced excitation of CSN was sustained and lasted the whole experiment (46 ± 7%, 55 ± 8%, 60 ± 10% at 60, 90, and 120 minutes after TNF-α administration, respectively; Figure 2B). It is important to highlight that throughout the experimental protocol, the animals were artificially ventilated with a slight hyperoxia (50% O_2_, balance N_2_) avoiding any potential hypoxia episode. We also performed additional experiments which demonstrated that the intravenous TNF-α did not affect the partial pressure of oxygen (PaO_2_), the partial pressure of carbon dioxide (PaCO_2_), the pH, and the bicarbonate (HCO_3_^−^) concentration in the arterial blood of unanesthetized, spontaneously breathing rats, confirming that the treatment does not produce hypoxia, hypercapnia or acidosis. (figure supplement 1). Thus, our data indicate that TNF-α can stimulate the carotid body and increase CSN activity independently of changes in blood gases and pH alterations. We, therefore, hypothesized that this TNF-α-induced increase in CSN afferent activity could activate central pathways similar to those activated by hypoxic stimuli, generating autonomic responses such as the activation of the sympathetic nervous system.

**Figure 2.**
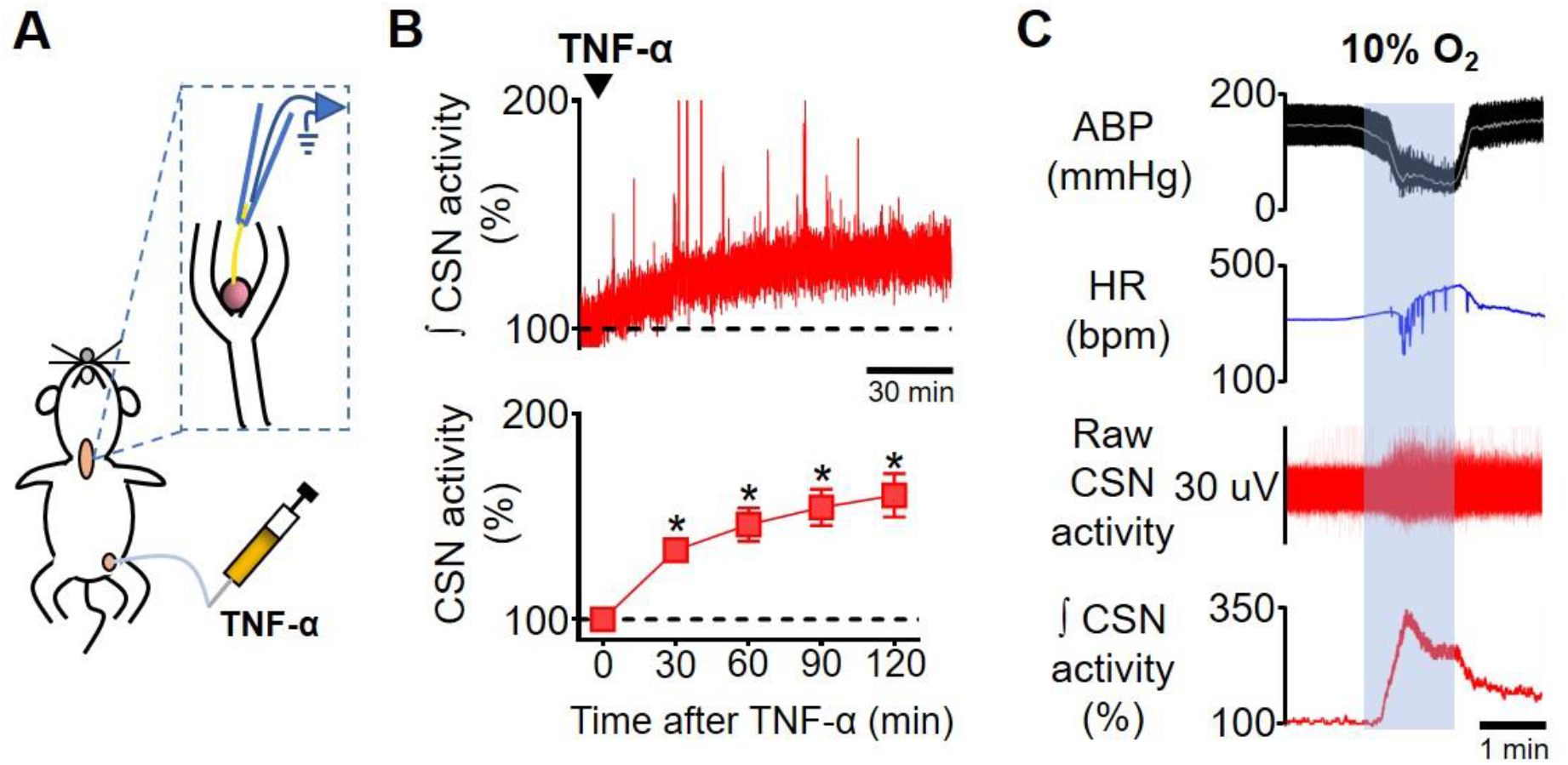
Carotid sinus nerve afferent activity (CSN activity) in response to intravenous TNF-α. **A.** Schematic illustration of the experimental protocol. **B.** Representative trace of the integrated CSN activity (∫ CSN activity; time constant = 1 s) from one rat during baseline and after TNF-α (500 ng, IV, black arrowhead) administration (top; scale bar = 30 minutes) and summary data showing CSN activity at baseline and 30, 60, 90 and 120 minutes after TNF-α administration (bottom; n = 6). Baseline CSN activity was normalized to 100% after noise subtraction. A one-way repeated measures ANOVA detected statistically significant differences in CSN activity over time, *F*_(4, 20)_ = 21,282, p < 0.001. Subsequent post hoc analysis with a Bonferroni adjustment revealed that, as compared to time 0 (baseline), CSN activity was statistically significantly higher at 30 minutes (34%, 95% CI [9, 59], p = 0.014); at 60 minutes (46%, 95% CI [8, 85], p = 0.023); at 90 minutes (55%, 95% CI [13, 96], p = 0.016); and at 120 minutes (60%, 95% CI [10, 111], p = 0.023) after TNF-α administration. *p < 0.05. Data are means ± SEM. **C.** Representative traces showing the viability of CSN activity recordings assessed by a brief exposure to hypoxia (10% O_2_, balance N_2_; grey shaded area). The typical acute response of urethane-anesthetized rats to hypoxia includes hypotension, bradycardia, and a robust increase in CSN activity. ABP, arterial blood pressure; HR, heart rate.

### RVLM-projecting cNTS glutamatergic neurons are activated by TNF-α

The first synapse of carotid body afferents within the central nervous system occurs in the cNTS, as extensively described in the literature (Colombari et al., 1996; Cruz et al., 2010; Kline et al., 2010; Malheiros-Lima et al., 2020). The cNTS sends excitatory glutamatergic projections to several areas, being implicated in diverse physiological functions. In the context of the carotid body-related functions, the cNTS neurons project to important autonomic areas involved in the neural control of cardiovascular and respiratory functions (Kline et al., 2010; Zera et al., 2019). For example, a previous report demonstrated direct monosynaptic projections from cNTS to RVLM, where the majority of pre-sympathetic neurons are located (Kline et al., 2010). It was also shown that most of these RVLM-projecting cNTS neurons are activated by hypoxia and constitute the major neural pathway of hypoxia-induced sympathetic activation (Kline et al., 2010; Koshiya and Guyenet, 1996). Thus, we sought to investigate if this sympathoexcitatory pathway is activated by circulating TNF-α since this cytokine increased the discharge of carotid body afferents, as shown in Figure 2B. Our results demonstrated massive monosynaptic projections from cNTS to RVLM (FG-labeled cells, blue staining, Figure 3C – E) in both SHAM and CB-X rats at all evaluated rostro-caudal levels: −14.40 mm to −14.64 mm (SHAM, 42 ± 7 cells; CB-X, 41 ± 7 cells), − 14.16 mm to −14.40 mm (SHAM, 43 ± 8 cells; CB-X, 36 ± 4 cells), and −13.92 mm to − 14.16 mm. (SHAM, 53 ± 6 cells; CB-X, 50 ± 8 cells). Most of these projections are excitatory (VGluT2^+^ cells, green staining, Figure 3C – E). Circulating TNF-α activated a considerable proportion of these RVLM-projecting glutamatergic cNTS neurons in SHAM rats, as indicated by c-FOS expression (red staining) in FG^+^/VGluT2^+^ cells (Figure 3C – E); Importantly, the number of activated RVLM-projecting glutamatergic cNTS neurons was dramatically reduced by carotid body ablation: −14.40 mm to −14.64 mm (SHAM, 11 ± 1 cells; CB-X, 2 ± 1 cells), −14.16 mm to −14.40 mm (SHAM, 17 ± 7 cells; CB-X, 3 ± 1 cells), and −13.92 mm to −14.16 mm. (SHAM, 18 ± 4 cells; CB-X, 3 ± 1 cells). The efficacy of the bilateral carotid body ablation procedure was confirmed by the lack of cardiovascular responses to KCN (figure supplement 2A – B). Together with our previous findings (Figures 1 and 2), these results suggest that the carotid body detects the circulating TNF-α through TNFR1 and transmits this information to the central nervous system via carotid sinus nerve afferents, resulting in the activation of a sympathoexcitatory pathway.

**Figure 3.**
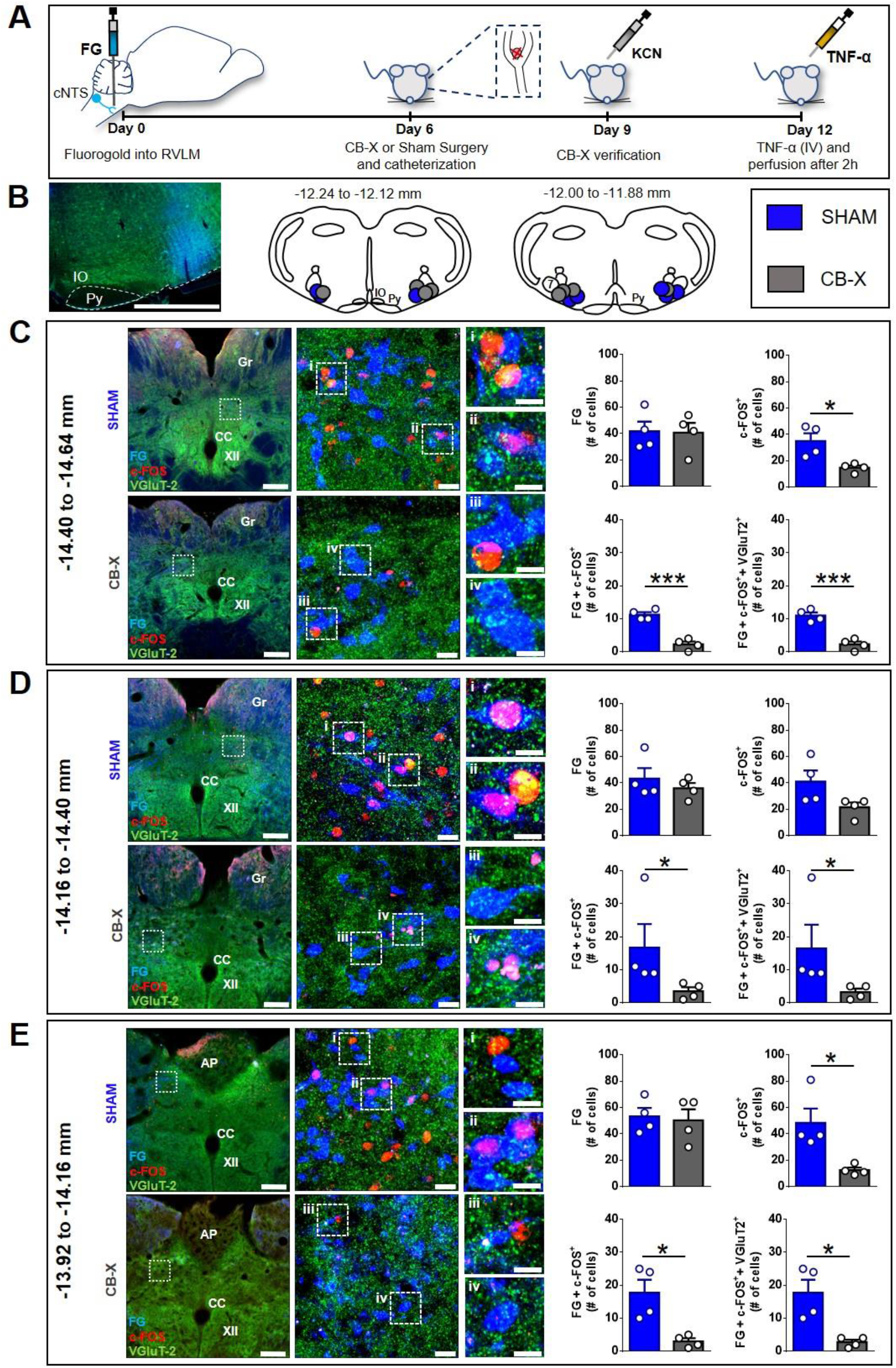
Activation of RVLM-projecting cNTS glutamatergic neurons by circulating TNF-α in SHAM and CB-X rats. **A.** Schematic illustration of the experimental protocol. **B.** Representative image from a typical retrograde tracer (Fluorogold; FG) injection-site into RVLM and schematic pictures of RVLM injections-sites of all bilaterally FG-injected animals (n=4 per group). IO, inferior olive; Py, pyramidal tract; 7, facial motor nucleus. Scale bar is 1000 µm. **C, D** and **E.** Images are representative pictures of cNTS sections at three different rostro-caudal levels, processed for c-FOS (red) and VGluT2 (green) immunofluorescence, and containing FG-positive cells retrogradely labeled from the RVLM (blue). Gr, gracile nucleus; CC, central canal; XII, hypoglossal nucleus; AP, area postrema. Scale bars are 200 µm for 5x magnification pictures (left), 20 µm for 40x magnification pictures (middle) and 10 µm for zoom pictures (right). **i, ii, iii**, and **iv.** Digital zoom into selected regions. Bar graphs show the quantification of retrogradely labeled FG neurons, c-FOS^+^ neurons, double stained (FG/c-FOS^+^) neurons and triple stained (FG/c-FOS^+^/VGluT2^+^) neurons in the cNTS 2 hours after TNF-α administration (500 ng, IV) in SHAM (n=4) and CB-X (n=4) rats. The number of RVLM-projecting neurons (FG-labeled cells) was not different between SHAM and CB-X rats in all evaluated cNTS levels: −14.40 to −14.64 mm (C), *t*(6) = 0.096, p = 0.926 (Student’s *t*-test); −14.16 to −14.40 mm (D), *U* = 7.5, Z = −0.145, p = 0.886 (Mann-Whitney *U*-test); and −13.92 to −14.16 mm (E), *t*(6) = 0.285, p = 0.785 (Student’s *t*-test). General neuronal activation (i.e., both RVLM-projecting and RVLM- non-projecting; c-FOS^+^ cells) was higher in SHAM as compared to CB-X rats at −14.40 to −14.64 mm (C), *t*(3.505) = 3.326, p = 0.036 (Welch’s *t*-test) and at −13.92 to −14.16 mm (E), *U* = 0, _Z_ = −2.323, p = 0.029 (Mann-Whitney *U*-test); but not at −14.16 to −14.40 mm (D), *t*(6) = 2.141, p = 0.076 (Student’s *t*-test). The specific activation of RVLM-projecting neurons (c-FOS^+^/FG^+^ cells) was higher in SHAM as compared to CB-X rats in all 3 cNTS levels: −14.40 to −14.64 mm (C), *t*(6) = 7.919, p < 0.001 (Student’s *t*-test); −14.16 to −14.40 mm (D), *U* = 0, _Z_ = −2.323, p = 0.029 (Mann-Whitney *U*-test); and −13.92 to −14.16 mm (E), *t*(3.324) = 3.661, p = 0.030 (Welch’s *t*-test). Virtually all activated RVLM-projecting cNTS neurons are glutamatergic (FOS^+^/FG/VGluT2^+^ cells). The number of activated RVLM-projecting cNTS glutamatergic neurons was higher in SHAM as compared to CB-X rats in all 3 cNTS levels: −14.40 to −14.64 mm (C), *t*(6) = 7.000, p < 0.001 (Student’s *t*-test); −14.16 to −14.40 mm (D), *U* = 0, _Z_ = −2.337, p = 0.029 (Mann-Whitney *U*-test); and −13.92 to −14.16 mm (E), *t*(3.219) = 3.755, p = 0.029 (Welch’s *t*-test). *p < 0.05 and ***p < 0.001. Data are means ± SEM.

### TNF-a promotes a carotid-body mediated increase in splanchnic SNA

Because circulating TNF-α activated a well-known sympathoexcitatory central pathway, we next performed experiments to investigate the effect of this cytokine on sympathetic activity directly recorded from multiple sympathetic nerves in vivo. Our results showed that intravenously administered TNF-α promotes a generalized sympathoexcitation in SHAM rats (Figure 4B – F), consistent with the activation of the RVLM-projecting cNTS glutamatergic neurons demonstrated in Figure 3: Δ Splanchnic SNA (14 ± 4%, 25 ± 7%, 32 ± 9% and 42 ± 11% respectively at 30, 60, 90, and 120 minutes after TNF-α administration); Δ Renal SNA (9 ± 4%, 18 ± 6%, 22 ± 8%, 27 ± 9% respectively at 30, 60, 90, and 120 minutes after TNF-α administration) and Δ lumbar SNA (5 ± 1%, 11 ± 3%, 16 ± 4%, 19 ± 5% respectively at 30, 60, 90, and 120 minutes after TNF-α administration). Interestingly, despite the generalized sympathetic activation, mean arterial blood pressure (ABP) only slightly increased (Figure 4B – C).

**Figure 4.**
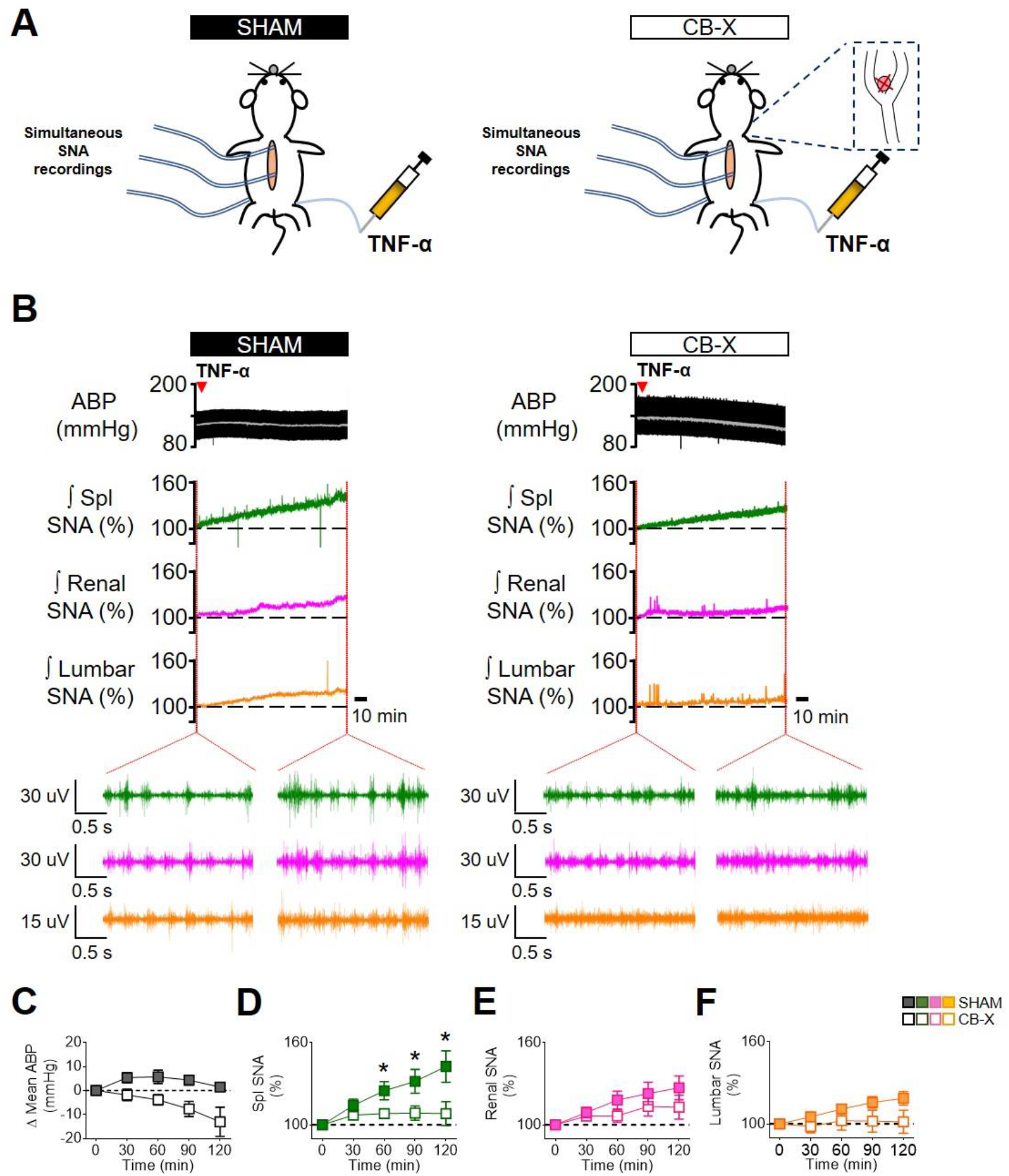
Carotid body ablation attenuates the TNF-α-induced splanchnic sympathetic activation. **A.** Schematic illustration of the experimental protocol. **B.** Representative traces of arterial blood pressure (pulsatile ABP, black; mean ABP, white), splanchnic (Spl; green), renal (magenta) and lumbar (orange) integrated (∫; time constant = 1s) sympathetic nerve activity (SNA) in sham-operated rats (SHAM) and carotid body-ablated rats (CB-X) during baseline conditions and in the next 2 hours after TNF-α administration (500 ng, IV, red arrowhead). For each sympathetic nerve, raw SNA signals at baseline and 2 hours after TNF-α administration are also presented (as indicated by the red dotted lines). **C, D, E and F.** Summary data showing the changes in mean ABP (C), Spl SNA (D), Renal SNA (E) and Lumbar SNA (F) in response to TNF-α in SHAM (filled symbols, n = 6) and CB-X (open symbols, n = 6) rats. For each rat, baseline integrated SNA was normalized to 100%, and the relative changes were calculated at four different time points (30, 60, 90 and 120 minutes after TNF-α administration). A statistically significant group x time interaction on spl SNA was detected by two-way repeated-measures ANOVA, *F*_(3,15)_ = 11.119, p < 0.001. Subsequent simple main effects analyses revealed that spl SNA changes were significantly greater in SHAM as compared to CB-X rats at 60 minutes, *F*_(1,5)_ = 7.042, p = 0.045; at 90 minutes, *F*_(1,5)_ = 10.224, p = 0.024; and at 120 minutes, *F*_(1,5)_ = 16.515, p = 0.010 after TNF-α administration. *p < 0.05. There were no statistically significant group x time interactions on Mean ABP, *F*_(1.113, 5.567)_ = 0.807, p = 0.420, ε = 0.371; Renal SNA, *F*_(3,15)_ = 0.805, p = 0.510; and Lumbar SNA, *F*_(3,15)_ = 1.685, p = 0.213 (two-way repeated measures ANOVA). Data are means ± SEM.

Since carotid body ablation almost abolished the TNF-α-induced activation of RLVM-projecting cNTS glutamatergic neurons (Figure 3), we tested whether the carotid bodies would be necessary to the observed sympathoexcitation in response to TNF-α administration. To accomplish that, we administered TNF-α to rats subjected to bilateral carotid body ablation (Figure 4B – F). CB-X rats displayed an attenuated increase in SNA in response to TNF-α: Δ Splanchnic SNA (7 ± 1%, 8 ± 2%, 8 ± 5% and 8 ± 9% respectively at 30, 60, 90, and 120 minutes after TNF-α administration), Δ renal SNA (6 ± 3%, 6 ± 5%, 13 ± 7% and 13 ± 9% respectively at 30, 60, 90, and 120 minutes after TNF-α administration), and Δ lumbar SNA (−2 ± 5%, 2 ± 7%, 2 ± 8% and 1 ± 8% respectively at 30, 60, 90, and 120 minutes after TNF-α administration). These SNA responses were diminished compared to those displayed by SHAM rats, especially on splanchnic SNA at 60, 90, and 120 minutes after TNF-α administration, suggesting that the carotid bodies contribute to this specific response (Figure 4D). Unlike SHAM rats, mean ABP in CB-X rats tended to decrease even without reductions in the activity of any of the recorded sympathetic nerves (Figure 4B – C). At the end of the experiments, bilateral carotid body ablation was confirmed by the lack of sympathetic and blood pressure responses to KCN (figure supplement 3A – B).

### Carotid body ablation or splanchnic sympathetic denervation exacerbates TNF-α-induced inflammation

Considering that the exogenous TNF-α activated a carotid body-cNTS-RVLM circuitry to excite a specific sympathetic nerve (splanchnic), and because the splanchnic sympathetic nerves have been considered essential components of sympathetic-mediated mechanisms to control inflammation (Lankadeva et al., 2020; Martelli et al., 2014), we next investigated if the activation of this newly described circuit could play an anti-inflammatory role in the TNF-α-induced inflammation. We found that, in SHAM rats that received TNF-α, the plasma levels of this cytokine (8.5 ± 1.3 pg mL^−1^) were found significantly higher in comparison to SHAM rats that received vehicle (1.1 ± 0.2 pg mL^−1^) (Figure 5B). It is important to mention that, the half-life of TNF-α is very short (few minutes) (Ma et al., 2015; Simó et al., 2012), and, hence, it is very likely that the measured levels of this cytokine in the plasma (2 hours after TNF-α or vehicle administrations) reflect endogenously produced TNF-α. In rats subjected to either carotid body ablation (CB-X) or splanchnic sympathetic denervation (SPL-X), the administration of TNF-α resulted in significant higher plasma levels of this cytokine compared to SHAM rats injected with TNF-α (CB-X + TNF-α = 58.1 ± 4.7 pg mL^−1^, SPL-X + TNF-α = 54.8 ± 11.8 pg mL^−1^) (Figure 5B), suggesting that the absence of the carotid bodies or the splanchnic sympathetic nerves exacerbated the systemic inflammatory status triggered by the exogenous TNF-α. In the same direction, the levels of TNF-α in the spleen were found higher in CB-X + TNF-α (4.5 ± 0.6 pg mg^−1^) and in SPL-X + TNF-α (5.1 ± 0.8 pg mg^−1^) groups compared to SHAM + TNF-α (1.7 ± 0.2 pg mg^−1^) group (Figure 5E). These results support the idea that the exogenously administered TNF-α induced the endogenous production of additional TNF-α likely via stimulation of splenic macrophages and, that, the removal of the carotid bodies (a potential sensor of TNF-α) or of the splanchnic sympathetic nerves (a potential suppressor of spleen-derived TNF-α production), significantly increased TNF-α levels in the spleen. It is important to highlight that in SPL-X + vehicle animals, the levels of TNF-α in the spleen were also elevated (6.0 ± 1.3 pg mg^−1^) (Figure 5E), reinforcing the notion that the splanchnic sympathetic innervation of the spleen (via celiac ganglion), exerts a kind of inhibitory tonus over splenic production of TNF-α. By way of comparison, in rats with intact splanchnic nerves (SHAM and CB-X) injected with vehicle, the levels of TNF-α in the spleen were low: (SHAM + vehicle = 1.0 ± 0.1 pg mg^−1^, CB-X + vehicle = 1.2 ± 0.1 pg mg^−1^) (Figure 5E).

**Figure 5.**
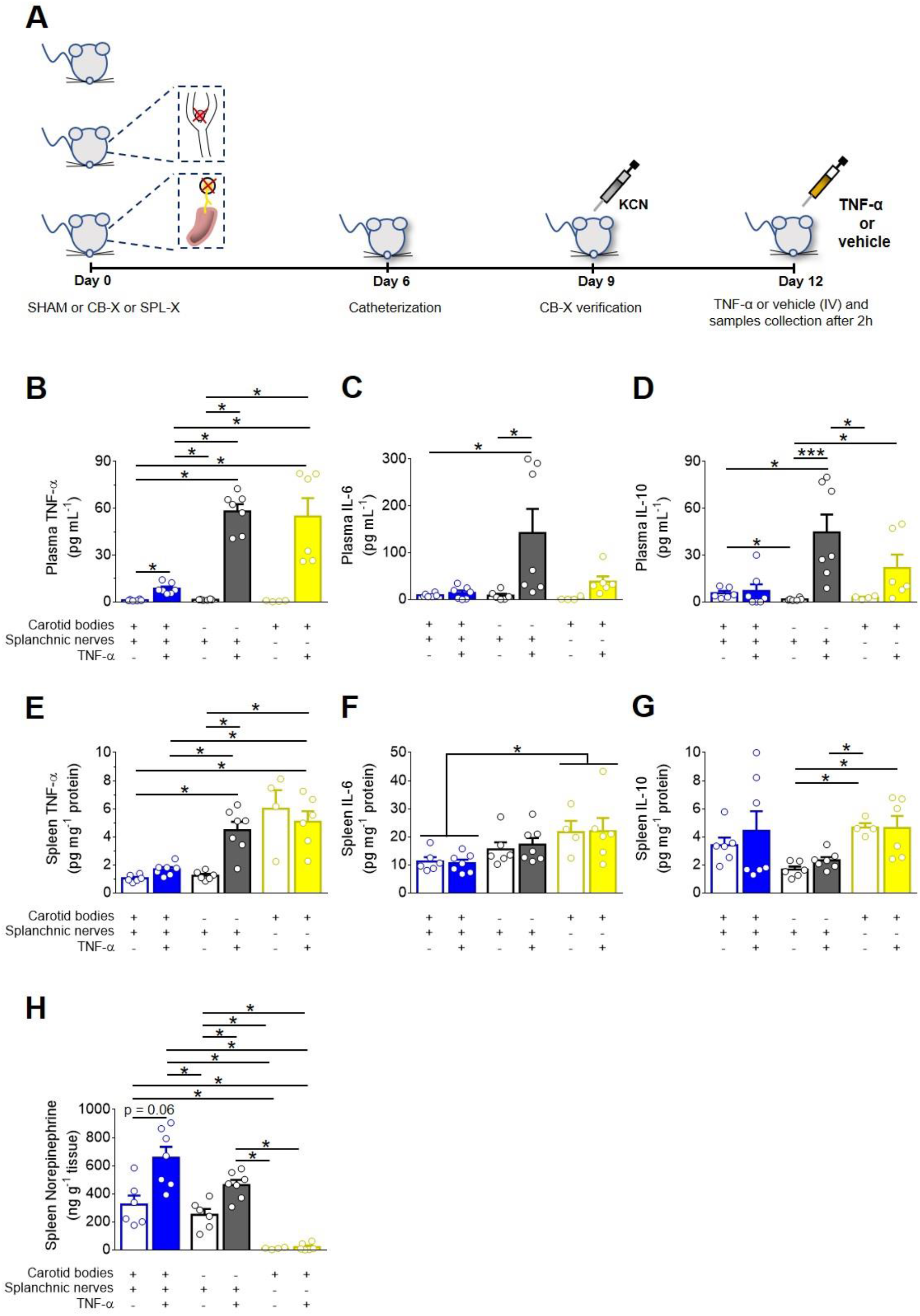
Carotid body ablation (CB-X) or splanchnic sympathetic denervation (SPL-X) intensify the TNF-α-induced inflammation. **A.** Schematic illustration of the experimental protocol. **B, C and D.** Plasma levels of TNF-α, IL-6 and IL-10 in SHAM (blue bars), CB-X (gray bars), and SPL-X (yellow bars) rats, measured 2 hours after vehicle (empty bars) or TNF-α (filled bars) intravenous administration (n = 4 - 7 per group). **B.** Statistically significant differences in the plasma levels of TNF-α across groups were detected: *H*(5) = 31.454, p < 0.001 (Kruskal-Wallis). In SHAM + TNF-α, the plasma levels of this cytokine were found significantly higher in comparison to SHAM + vehicle (*U* = 0, _Z_ = −3.000, p = 0.001, Mann-Whitney *U*-test). In CB-X and SPL-X rats, TNF-α administration resulted in significant higher plasma levels of this cytokine as compared to SHAM + TNF-α: SHAM + TNF-α vs. CB-X + TNF-α (*U* = 0, _Z_ = - 3.130, p = 0.001, Mann-Whitney *U*-test); SHAM + TNF-α vs. SPL-X + TNF-α (*U* = 0, _Z_ = −3.000, p = 0.001, Mann-Whitney *U*-test). Between vehicle-treated groups, the plasma levels of TNF-α were not different (p > 0.003). Regarding plasma IL-6 levels, significant differences between groups were detected: *H*(5) = 22.024, p = 0.001 (Kruskal-Wallis). **C.** The plasma levels of IL-6 were higher in CB-X + TNF-α as compared to SHAM + vehicle (*U* = 1, _Z_ = −2.857, p = 0.002, Mann-Whitney *U*-test) and to CB-X + vehicle (*U* = 1, _Z_ = −2.857, p = 0.002, Mann-Whitney *U*-test). No statistical differences were found in plasma levels of IL-6 in the other pairwise comparisons (p > 0.003). **D.** Finally, the plasma levels of IL-10 were found significantly different across groups: *F*_(5, 13.522)_ = 14.524, p < 0.001 (Welch ANOVA). Games-Howell post hoc test revealed that the plasma levels of IL-10 were significantly higher in CB-X + TNF-α as compared to all groups that received vehicle: CB-X + TNF-α vs. SHAM + vehicle (Mean difference = 2.0 pg mL^−1^, 95% CI [0.6, 3.3], p = 0.005); CB-X + TNF-α vs. CB-X + vehicle (Mean difference = 3.1 pg mL^−1^, 95% CI [1.8, 4.4], p < 0.001); CB-X + TNF-α vs. SPL-X + vehicle (Mean difference = 2.6 pg mL^−1^, 95% CI [1.2, 3.9], p = 0.001). In SPL-X + TNF-α, the plasma levels of IL-10 were higher as compared to CB-X + vehicle (Mean difference = 2.2 pg mL^−1^, 95% CI [0.2, 4.1], p = 0.032). Between vehicle-administered groups, SHAM rats displayed higher plasma levels of IL10 as compared to CB-X rats (Mean difference = 1.2 pg mL^−1^, 95% CI [0.1, 2.2], p = 0.033. **E, F and G.** Spleen levels of TNF-α, IL-6 and IL-10 in SHAM (blue bars), CB-X (gray bars), and SPL-X (yellow bars) rats, 2 hours after vehicle (empty bars) or TNF-α (filled bars) intravenous administration (n = 4 - 7 per group). **E.** Statistically significant differences in the spleen levels of TNF-α between groups were found: *F*_(5, 12.262)_ = 12.957, p < 0.001 (Welch ANOVA). Games-Howell post hoc test revealed that the spleen levels of TNF-α were significantly higher in CB-X and SPL-X that received TNF-α as compared to SHAM rats that received TNF-α: CB-X + TNF-α vs. SHAM + TNF-α (Mean difference = 2.8 pg mg^−1^ protein, 95% CI [0.5, 5.1], p = 0.021); SPL-X + TNF-α vs. SHAM + TNF-α (Mean difference = 3.4 pg mg^−1^ protein, 95% CI [0.2, 6.6], p = 0.039). Within vehicle-treated groups, the spleen levels of TNF-α were not different (p > 0.05). No statistical differences were found when comparing SHAM + vehicle vs. SHAM + TNF-α (Mean difference = −0.6 pg mg^−1^ protein, 95% CI [−1.3, 0.0], p = 0.064). **F.** Regarding the spleen levels of IL-6, no interactions between group x treatment were detected: *F*_(2,30)_ = 0.092, p = 0.912, partial η^2^ = 0.006. However, a statistically significant main effect of group was found: *F*_(2,30)_ = 7.130, p = 0.003, partial η^2^ = 0.322. A Bonferroni post hoc analysis indicated that the spleen levels of IL-6 were significant higher in SPL-X groups as compared to SHAM groups: (Mean difference = 10.9 pg mg protein, 95% CI [3.5, 18.3], p = 0.002. **G.** Concerning the spleen levels of IL-10, statistically significant differences between groups were found: *F*_(5, 13.792)_ = 12.491, p < 0.001 (Welch ANOVA). Games-Howell post hoc test revealed that the spleen levels of IL-10 were significantly lower in CB-X groups as compared to SPL-X groups: CB-X + vehicle vs. SPL-X + vehicle (Mean difference = −1.1 pg mg^−1^ protein, 95% CI [− 1.6, −0.5], p = 0.002); CB-X + vehicle vs. SPL-X + TNF-α (Mean difference = −1.0 pg mg^−1^ protein, 95% CI [−1.8, −0.1], p = 0.026); CB-X + TNF-α vs. SPL-X + vehicle (Mean difference = −0.7 pg mg^−1^ protein, 95% CI [−1.1, −0.3], p = 0.002). **H.** Spleen levels of norepinephrine in SHAM (blue bars), CB-X (gray bars), and SPL-X (yellow bars) rats, 2 hours after vehicle (empty bars) or TNF-α (filled bars) intravenous administration (n = 4 - 7 per group).. Statistically significant differences in the spleen levels of norepinephrine between groups were found: *F*_(5, 13.050)_ = 45.864, p < 0.001 (Welch ANOVA). *p < 0.05 and ***p < 0.001. Data are means ± SEM. Games-Howell post hoc test revealed that the administration of TNF-α in SHAM rats, resulted in a trend to increase the spleen norepinephrine levels compared to SHAM animals receiving vehicle (Mean difference = 333.3 pg mg^−1^ tissue, 95% CI [−11.2, 677.8], p = 0.060) and in significant increases as compared to CB-X + vehicle (Mean difference = 406.3 pg mg^−1^ tissue, 95% CI [94.4, 718.2], p = 0.011), to SPL-X + vehicle (Mean difference = 645.9 pg mg^−1^ tissue, 95% CI [337.1, 954.7], p = 0.001), and to SPL-X + TNF-α (Mean difference = 635.5 pg mg^−1^ tissue, 95% CI [327.6, 943.5], p = 0.001). In CB-X rats, TNF-α administration led to higher levels of norepinephrine in the spleen as compared to CB-X + vehicle (Mean difference = 212.4 pg mg^−1^ tissue, 95% CI [24.4, 400.4], p = 0.025), to SPL-X + vehicle (Mean difference = 452.0 pg mg^−1^ tissue, 95% CI [307.5, 596.6], p < 0.001), and to SPL-X + TNF-α (Mean difference = 441.7 pg mg^−1^ tissue, 95% CI [298.1, 585.3], p < 0.001). SPL-X + vehicle animals also displayed lower levels of norepinephrine in the spleen compared to SHAM + vehicle (Mean difference = −312.6 pg mg^−1^ tissue, 95% CI [−587.0, −38.1], p = 0.030) and CB-X + vehicle (Mean difference = −239.6 pg mg^−1^ tissue, 95% CI [−413.5, −65.7], p = 0.013). Similarly, the levels of norepinephrine in the spleen were also lower in SPL-X + TNF-α compared to SHAM + vehicle (Mean difference = −302.2 pg mg^−1^ tissue, 95% CI [−574.7, −29.8], p = 0.033) and CB-X + vehicle (Mean difference = −229.3 pg mg^−1^ tissue, 95% CI [−400.7, −57.9], p = 0.014).

**Figure 6.**
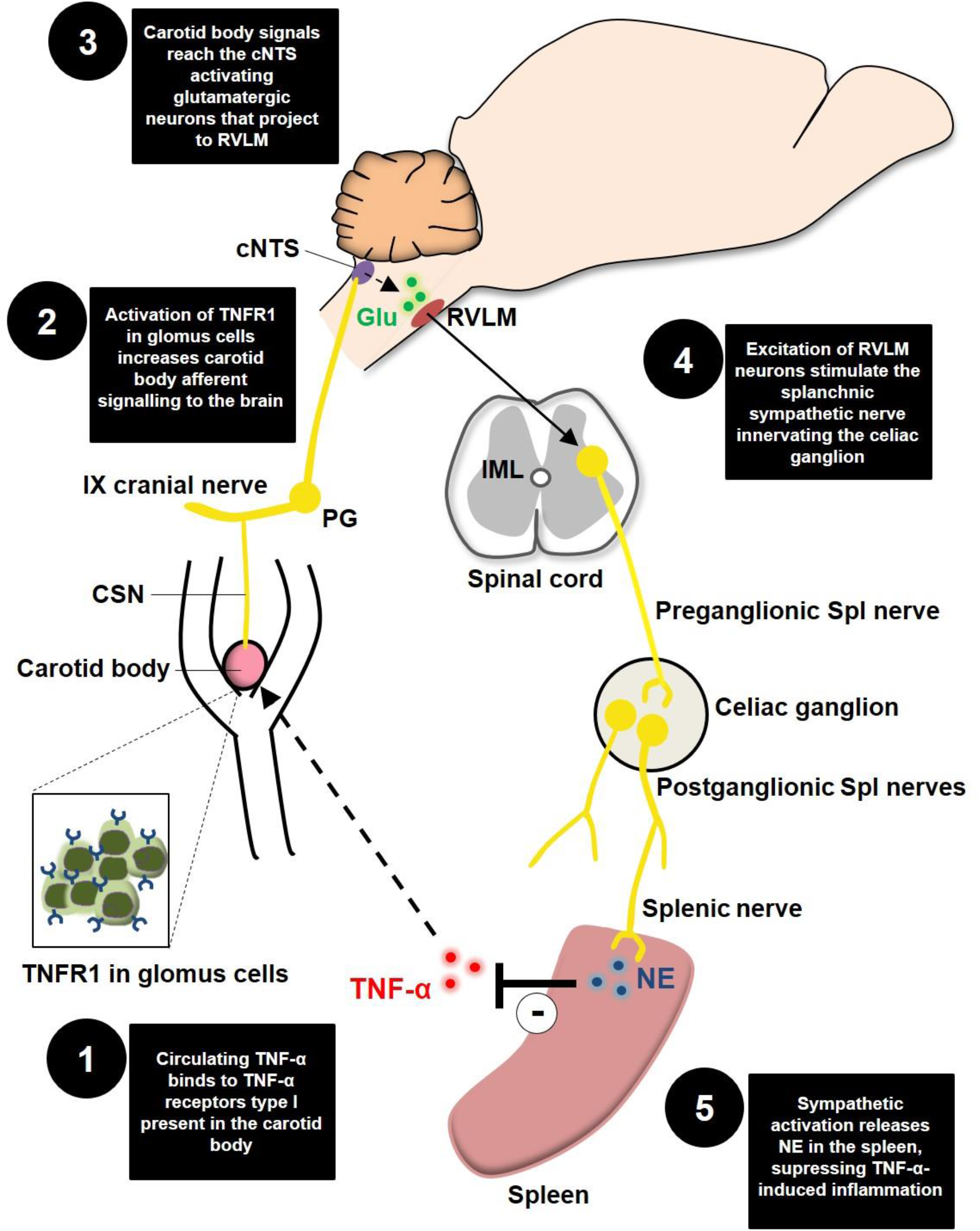
Schematic model of the novel proposed neuroimmune mechanism. TNFR1, TNF-α receptors type I; CSN, carotid sinus nerve; IX cranial nerve, glossopharyngeal nerve; PG, petrosal ganglion; cNTS, commissural nucleus tractus solitarius; Glu, glutamate; RLVM, rostral ventrolateral medulla; IML, intermediolateral nucleus; Spl, splanchnic; NE, norepinephrine.

Regarding plasma IL-6 levels, CB-X + TNF-α animals displayed higher levels (142.2 ± 51.0 pg mL^−1^) than SHAM + vehicle (9.1 ± 1.6 pg mL^−1^) and CB-X + vehicle (8.2 ± 3.8 pg mL^−1^) (Figure 5C). Although not statistically significant, the levels of IL-6 in the plasma tended to be higher in CB-X + TNF-α and SPL-X + TNF-α (38.1 ± 11.2 pg mL^−1^) compared to all other groups: (SHAM + vehicle = 9.1 ± 1.6 pg mL^−1^, SHAM + TNF-α = 14.1 ± 5.0 pg mL^−1^, CB-X + vehicle = 8.2 ± 3.8 pg mL^−1^, SPL-X + vehicle = 2.2 ± 1.9 pg mL^−1^) (Figure 5C). Concerning the spleen levels of IL-6, no interactions between group x treatment were detected by two-way ANOVA. However, a statistically main effect of group indicated that the spleen levels of IL-6 were higher in SPL-X + vehicle (21.7 ± 4.0 pg mg^−1^) and SPL-X + TNF-α (22.0 ± 4.7 pg mg^−1^) groups compared to SHAM + vehicle (11.2 ± 1.5 pg mg^−1^) and SHAM + TNF-α (10.6 ± 1.3 pg mg^−1^) groups (Figure 5F), suggesting that splanchnic sympathetic denervation was permissive to IL-6 production in the spleen, even in the absence of the TNF-α stimulus. With regard to plasma IL-10, the levels of this anti-inflammatory cytokine tended to be higher in CB-X + TNF-α (44.7 ± 11.4 pg mL^−1^) and SPL-X + TNF-α (21.7 ± 8.6 pg mL^−1^) as compared to SHAM + TNF-α (7.1 ± 4.2 pg mL^−1^) and to every other group that received vehicle (SHAM + vehicle = 5.6 ± 1.4 pg mL^−1^, CB-X + vehicle = 1.7 ± 0.3 pg mL^−1^, SPL-X + vehicle = 2.8 ± 0.5 pg mL^−1^) (Figure 5D). These results match with the increased levels of TNF-α in the plasma and the spleen of CB-X and SPL-X rats that received TNF-α, indicating a worse systemic inflammatory status in these animals. Finally, spleen levels of IL-10 tended to be lower in both CB-X groups, but was only statistically different between: CB-X + vehicle (1.7 ± 0.2 pg mg^−1^) compared to SPL-X + vehicle (4.7 ± 0.3 pg mg^−1^); CB-X + vehicle group compared to SPL-X + TNF-α (4.6 ± 0.9 pg mg^−1^); and between CB-X + TNF-α (2.3 ± 0.2 pg mg^−1^) compared to SPL-X + vehicle (Figure 5G). In regard to norepinephrine levels in the spleen, SHAM rats injected with TNF-α displayed the highest mean levels (656.2 ± 77.6 pg mg^−1^), followed by CB-X + TNF-α (462.3 ± 36.4 pg mg^−1^), SHAM + vehicle (322.9 ± 64.4 pg mg^−1^), CB-X+ vehicle (249.9 ± 40.9 pg mg^−1^), SPL-X + TNF-α (20.6 ± 10.7 pg mg^−1^), and SPL-X + vehicle (10.3 ± 3.5 pg mg^−1^) groups (Figure 5H). Note that splanchnic sympathetic denervation almost depleted the norepinephrine content in the spleen, confirming the efficacy of the denervation procedure. In addition, the efficacy of splanchnic sympathetic denervation was also verified by the less pronounced TH staining in the spleen (figure supplement 4C). The efficacy of the bilateral carotid body ablation procedure was confirmed by the lack of cardiovascular responses to KCN (figure supplement 4A – B). Collectively, our data suggest that elevated circulating levels of TNF-α activates a neural mechanism (carotid body-cNTS-RVLM-splanchnic sympathetic nerves) that controls the ongoing inflammation by inhibiting the synthesis of additional TNF-α in the spleen likely via direct norepinephrine-mediated suppression of splenic macrophage TNF-α production.

## Discussion

In the present study, we provide a series of anatomical and functional evidence for the existence of a previously unrecognized mechanism of neuroimmune interaction. The main finding is that the carotid body is able to detect elevated levels of the pro-inflammatory cytokine TNF-α in the blood and communicate with the central nervous system via carotid sinus nerve afferents, activating RVLM-projecting cNTS excitatory neurons that contribute to a counteracting sympathetic-mediated anti-inflammatory response. These results advance our understanding of the complex mechanisms underlying the bidirectional connection between the nervous and the immune systems.

Recently, the carotid bodies emerged as potential candidates for peripheral detectors of inflammation. This possibility is supported by a growing number of studies indicating that they are polymodal sensors, able to monitor the chemical composition of the arterial blood. More specifically, these studies have shown that besides promoting autonomic and respiratory adjustments in response to arterial hypoxemia (i.e., peripheral chemoreflex), the carotid bodies can respond to several other circulating stimuli such as leptin, angiotensin II, glucose, sodium chloride, insulin, adrenaline, and, also, inflammatory mediators (Allen, 1998; da Silva et al., 2019; Jendzjowsky et al., 2021, 2018; Katayama, 2016; Kumar and Prabhakar, 2012; Shin et al., 2019; Thompson et al., 2016). Regarding inflammatory mediators, studies reported that the carotid body of many species, including rats, cats and humans, expresses receptors for lysophosphatidic acid (LPA), IL-1β, IL-6, and TNF-α (Fernández et al., 2008; Jendzjowsky et al., 2018; Mkrtchian et al., 2012; Wang et al., 2002). Accordingly, in the present study, we combined immunofluorescence and RNAscope FISH protocols to confirm that TNFR1 is expressed in the carotid body of rats at both mRNA and protein levels. Moreover, in addition to the anatomical evidence, previous functional studies demonstrated that inflammation-related factors can impact carotid body activity (Jendzjowsky et al., 2021, 2018; Shu et al., 2007), opening a wide range of possibilities regarding the role of the carotid body in the context of neuroimmune interactions. For instance, a recent study showed that LPA potently increased CSN activity in an isolated perfused carotid body/carotid sinus nerve preparation (Jendzjowsky et al., 2018). Furthermore, the same research group showed that the perfusion of the isolated carotid body/carotid sinus nerve preparation with diverse pro-inflammatory cytokines (IL-4, IL-5, IL-13, IL-1, IL-6, and TNF-α), one at a time or in combination, also increased CSN activity (Jendzjowsky et al., 2021), confirming the unique ability of the carotid body to sense and respond to inflammatory mediators. In our study, CSN activity was recorded in vivo and TNF-α was given systemically (IV). We chose the IV administration route because it better mimics a real scenario of systemic inflammation. We observed a progressive and significant increase in CSN activity, indicating that the carotid body could detect the elevated levels of TNF-α in the blood and alert the central nervous system via afferent signals. The reasons by which TNF-α increased CSN activity in a sustained manner (for at least 2 hours) are not clear, especially because the half-life of TNF-α in the plasma is reported to be very short (few minutes) (Ma et al., 2015; Simó et al., 2012). We hypothesize that the exogenous administered TNF-α stimulated the synthesis and release of additional TNF-α, probably via direct activation of splenic macrophages as suggested by our data (Figure 5) and/or by indirect activation of liver Kupffer cells as observed during endotoxemia in rats (Fonseca et al., 2021). This endogenously produced TNF-α could either sustain the carotid body activation and, also, stimulate the synthesis of further TNF-α.

We found that besides increasing CSN activity, the intravenous administration of TNF-α promoted the activation of cNTS neurons, the first relay site for carotid body afferents. Notably, the systemic administration of TNF-α resulted in activation of the cNTS neurons at the same rostro-caudal levels reported to be activated after carotid body stimulation by hypoxia or intravenous KCN (Cruz et al., 2010; Kline et al., 2010; Malheiros-Lima et al., 2020). These cNTS neurons, activated by carotid body stimulation, project to several brain areas, including the RVLM, to control the sympathetic nervous system (Kline et al., 2010; Koshiya and Guyenet, 1996). A previous study observed that after 3 hours of hypoxia (10% O_2_) exposure, a high proportion of RVLM-projecting cNTS neurons were activated (Kline et al., 2010). Furthermore, the authors injected anterograde tracers into the carotid body and observed that carotid body afferents terminate in close apposition to the RVLM-projecting cNTS neurons. Thus, this neural circuitry elegantly revealed by Kline et al. (2010), along with previous data (Aicher et al., 1996; Koshiya and Guyenet, 1996), provides a major neural pathway for hypoxia-induced sympathoexcitation. Of note, the blockade of glutamatergic receptors in the NTS was shown to strongly reduce the sympathetic responses to chemical stimulation of the carotid body (Ferreira et al., 2018). Since in the present study, circulating TNF-α induced the activation of RLVM-projecting cNTS glutamatergic neurons at the same rostro-caudal levels reported in the literature (Cruz et al., 2010; Kline et al., 2010; Malheiros-Lima et al., 2020) and, because carotid body ablation almost abolished the activation of these neurons, we believe that TNF-α might be stimulating a similar neural pathway (carotid body-cNTS-RVLM) activated by hypoxia to increase sympathetic activity. It is important to highlight that more than a half of c-FOS positive neurons observed in SHAM rats treated with TNF-α were not co-localized with FG (non-RVLM-projecting). We hypothesize that these neurons project to other nuclei involved in sympathetic modulation, such as the PVN, regions involved in respiratory control, and vagal nuclei (Luise King et al., 2012; Malheiros-Lima et al., 2020; Neff et al., 1998; Willis et al., 1996; Zera et al., 2019). In fact, recent studies suggested that inflammation-induced carotid body stimulation could also activate brainstem vagal nuclei (nucleus ambiguus and dorsal motor nucleus of the vagus) to increase parasympathetic activity (Jendzjowsky et al., 2021, 2018). Therefore, the results of the present and previous studies suggest that the carotid body detects circulating inflammatory mediators and activates central autonomic areas to modulate sympathetic and/or parasympathetic functions.

Our study shows that the TNF-α-induced activation of a sympathoexcitatory circuit (carotid body-cNTS-RVLM) resulted in increased SNA as revealed by simultaneous recordings of splanchnic, renal and lumbar SNA. To the best of our knowledge, this is the first study describing the effects of circulating TNF-α, an important inflammatory mediator, on the activity of three different sympathetic nerves recorded simultaneously in vivo. Previous studies have already demonstrated that circulating TNF-α increases renal SNA in rats (Wei et al., 2013; Zhang et al., 2003). However, since sympathetic outflows to other tissues/organs have distinct functions and can be differentially regulated (Morrison, 2001; Tromp et al., 2018), it becomes relevant to study the effects of TNF-α on sympathetic outflows directed to other targets besides the kidneys. Here, we found that TNF-α promoted a generalized activation of the sympathetic nervous system, increasing splanchnic, renal, and lumbar SNA in carotid body-intact rats. The removal of the afferent inputs from the carotid bodies (by bilateral carotid body ablation) blunted, in part, this TNF-α-induced sympathetic activation, consistent with the attenuated activation of RLVM-projecting cNTS neurons observed in CB-X rats (Figure 3C – E). Interestingly, the blunting effect of carotid body ablation was significant only on splanchnic SNA. Therefore, our data indicate that increased circulating TNF-α activates a carotid body-cNTS-RVLM neural circuit that selectively controls splanchnic SNA in this condition. It is noteworthy that a previous study reported that the increase in renal SNA following the systemic administration of TNF-α was largely attenuated in rats with lesions of the subfornical organ (Wei et al., 2013). It suggests that splanchnic, renal, and lumbar SNA might be under the control of different neural routes and might have different functions in the course of TNF-α-driven inflammation.

In this context, some studies have suggested that the splanchnic sympathetic nerves play an important immunomodulatory role during endotoxemia-induced systemic inflammation (Lankadeva et al., 2020; Martelli et al., 2014). For instance, it was demonstrated that acute endotoxemia induced by intravenous administration of lipopolysaccharide (LPS) significantly increased plasma levels of TNF-α after 90 minutes in rats (Martelli et al., 2014). In parallel, this LPS administration potently increased splanchnic SNA. Notably, when LPS was given to rats subjected to the bilateral section of the splanchnic sympathetic nerves, the plasma TNF-α levels increased 5 times more than those of intact rats (Martelli et al., 2014). Together, these results indicate that during LPS-induced systemic inflammation, the splanchnic SNA increases to counteract the ongoing inflammation in a kind of negative feedback reflex. Since, in the present study, the elevated circulating TNF-α activated a carotid body-cNTS-RVLM neural circuit to increase splanchnic SNA, we hypothesized that this mechanism could be a neuroimmune reflex to counteract the TNF-α-induced inflammation. To test this hypothesis, we removed either the detection/afferent arm (i.e., the carotid bodies) or the efferent arm (i.e., the splanchnic sympathetic nerves) of this potential neuroimmune reflex and subjected these animals (and SHAM control animals) to systemic injections of TNF-α or vehicle. After 2 hours, we quantified the levels of TNF-α, IL-6, and IL-10 in the blood and in the spleen as well as the levels of norepinephrine in the spleen. We found that in SHAM rats, the administration of TNF-α significantly increased the plasma levels of TNF-α and slightly increased the spleen levels of TNF-α compared to vehicle-injected SHAM rats. In addition, TNF-α administration tended to increase spleen norepinephrine levels in SHAM animals as compared to its vehicle-treated counterparts (Figure 5H, p = 0.06), consistent with our data showing a TNF-α induced splanchnic SNA activation. Interestingly, in rats subjected to either carotid body ablation or splanchnic sympathetic denervation, the administration of TNF-α resulted in exacerbated levels of pro-inflammatory cytokines in the plasma and the spleen, supporting the idea that both the detection/afferent arm and the efferent arm are important components of a neuroimmune regulatory mechanism that detects and modulates acute inflammation through sympathetic activation towards the spleen. Disrupting the afferent/detection component (carotid body ablation) resulted in a peculiar elevation of all quantified cytokines, including IL-10 (an anti-inflammatory cytokine). The reason for this elevation in plasma IL-10 in CB-X rats treated with TNF-α is not clear. This could result from the fact that carotid body ablation eliminated only part of the autonomic circuits toward the spleen, possible preserving and/or amplifying other counter-inflammatory mechanisms. In fact, the administration of TNF-α in CB-X rats, still activated splanchnic SNA and resulted in a significant increase in splenic levels of norepinephrine compared to vehicle-injected CB-X rats. However, the TNF-α-induced splanchnic SNA activation and norepinephrine release in the spleen were attenuated in CB-X rats compared to SHAM rats, which could explain, at least in part, the exacerbated inflammatory status observed in the animals lacking the carotid bodies. Differently, the interruption of the efferent component (splanchnic sympathetic denervation) completely blocked the sympathetic signalling to the spleen, removing the norepinephrine “inhibitory tonus” on cytokine production by splenic macrophages, resulting in elevated splenic cytokine levels even in those animals administered with saline. Collectively, our data suggest the existence of an intrinsic and physiological anti-inflammatory reflex that depends on a detection/afferent arm (i.e., the carotid bodies and the carotid sinus nerve), on a central integrative pathway (i.e., RVLM-projecting cNTS neurons), and on an effector/efferent arm (i.e., splanchnic sympathetic nerves) that modulates the splenic production of cytokines through norepinephrine release.

The findings of the present study are novel and place the carotid body as a critical player in the context of neuroimmune interactions. In the last years, the contribution of the carotid bodies to sympathetic overactivity has been implicated in the pathophysiology of several diseases such as sleep apnoea, hypertension, and heart failure (Marcus et al., 2014; McBryde et al., 2013; Melo et al., 2019; Narkiewicz et al., 2016; Niewinski et al., 2017; Yuan et al., 2016). In these conditions, exaggerated tonic CSN activity leads to chronic activation of the sympathetic nervous system, often associated with a poor prognosis. Here, we found that the acute carotid body-mediated sympathetic activation induced by intravenous TNF-α is likely to be beneficial because it exerted a counteracting anti-inflammatory reflex. However, it is possible that in chronic pathological inflammatory conditions, the long-term activation of this carotid body-dependent neuroimmune circuit leads to side effects because it generates an aberrant tonic CSN input to central sympathetic networks, leading to sustained sympathetic overactivity to multiple target organs. This possibility raises an intriguing question on whether circulating inflammatory factors could trigger the carotid body-mediated sympathetic overactivity observed in diseases such as hypertension (McBryde et al., 2013; Narkiewicz et al., 2016) and heart failure (Marcus et al., 2014; Niewinski et al., 2017) since these conditions are associated with increased systemic inflammation (Bautista et al., 2005; Norlander et al., 2018; Rauchhaus et al., 2000; Sesso et al., 2015). On the other hand, defects in the carotid body-mediated neuroimmune reflex described here, could impair the ability to regulate the levels of inflammatory mediators in the bloodstream, amplifying systemic inflammation. Nevertheless, further investigations are needed to clarify the beneficial or detrimental effects following the activation/inactivation of the neuroimmune mechanism described in the present study under different conditions and to explore its therapeutic potential in the treatment of inflammatory diseases.

## Methods

### Animals and ethical approval

All experimental procedures were reviewed and approved by the Ethical Committee in Animal Experimentation of the Araraquara School of Dentistry, São Paulo State University (protocol n° 17/2019) and conducted following the Guide for the Care and Use of Laboratory Animals from the Brazilian National Council for Animal Experimentation Control. Experiments were performed on adult male *Holtzman* rats (320 - 400 g) obtained from the Animal Care Unit of the São Paulo State University (Araraquara, SP, Brazil). The animals were housed in collective cages (2 - 4 animals/cage), provided with chow and water *ad libitum*, and maintained under controlled conditions of temperature (22 ± 1°C), humidity (50 - 60%) in a 12:12 hours light/dark cycle.

### General procedures

All surgical procedures were performed under aseptic conditions. The appropriate depth of anesthesia was confirmed by the absence of withdrawal reflex and corneal reflexes in response to pinching the toe. Throughout the surgical procedures and the experimental protocols performed under anesthesia (described below), the body temperature was measured by a rectal probe and maintained at 37 ± 0.5°C with a water-circulating heating pad.

### Experiment 1: Expression of TNF-α receptor type I in carotid body glomus cells

Rats were deeply anesthetized with isoflurane (5% in 100 O_2_) and subjected to transcardial perfusion with cold phosphate-buffered saline (PBS, 10 mM, pH 7.4, 100 mL/100 g BW) followed by paraformaldehyde (PFA, 4% in PBS, 100 mL/100 g BW). Whole carotid bifurcations containing the carotid bodies were collected as previously described (Pijacka et al., 2018) and fixed in PFA for 24 hours at 4° C. Next, carotid bifurcations were transferred to 10% sucrose solution and kept at 4° C until the tissue sinks. This procedure was repeated with 20% and 30% sucrose solutions. Carotid bifurcations were frozen in Tissue Freezing Medium (Triangle Biomedical Sciences, Durham, NC, USA) using dry ice, sectioned at 10 µm in a cryostat and mounted on microscope slides (Superfrost Plus, Fisher Scientific, Pittsburgh, PA, USA). To evaluate the expression of TNF-α receptor type I (TNFR1) in the carotid bodies, we employed two different approaches: 1) a fluorescent *in situ hybridization* (FISH) assay (RNAscope, Advanced Cell Diagnostics, Newark, CA, USA) for TNFR1 mRNA detection combined with immunofluorescence staining for TH (a marker of carotid body glomus cells) and; 2) a double immunofluorescence staining for TNFR1 and TH. In the first approach, the FISH assay was performed according to the manufacturer instructions (document #323100-USM, available at https://acdbio.com/documents/product-documents) and the following materials were used: RNAscope Multiplex Fluorescent Detection Reagents v2 (product #323110), the kit RNAscope H_2_O_2_ and Protease Reagents (product #322381), the RNAscope probe for TNFR1 (product #408111) and the TSA Cyanine 3 Plus Evaluation kit (product #NEL744001KT, Akoya Biosciences, Boston, MA, USA). After completing the FISH protocol, an immunofluorescence protocol for TH was performed to identify carotid body glomus cells. Briefly, the slides were incubated in a blocking solution (0.1 M PBS, 10% normal horse serum, and 0.3% Triton X-100) for 20 min and subsequently rinsed 3 x 10 minutes in 0.1 M PBS at room temperature. Then, the slides were incubated in primary antibody (Mouse anti-TH antibody, 1:1000, product #MAB5280, Millipore, Billerica, MA, USA) for 1 hour at room temperature and 36 hours at 4° C. After rinsing in PBS, the slides were incubated in secondary antibody (Alexa Fluor 488 donkey anti-mouse antibody, 1:200, product #R37114, Molecular Probes-Life Technologies, Eugene, OR, USA) for 4 hours at room temperature. The slides were rinsed in PBS, the excess liquid was drained, mounting medium (Fluoromount) was dropped on the tissue and slides were covered with glass coverslips (Fisherfinest). In the second approach, the immunofluorescence staining as carried out as described above but adding also a primary antibody for TNFR1 (Rabbit anti-TNFR1, 1:500, product #ab19139, Abcam, Cambridge, MA, USA) and a secondary antibody (Alexa Fluor 594 donkey anti-rabbit antibody, 1:200, product #A21207, Molecular Probes-Life Technologies). Images were acquired using a laser scanning confocal microscope (LSM800, Zeiss). For presentation purposes (color-blind safe) images were pseudo-colored and representative figures were prepared using the Zen 2 software (Blue edition, Zeiss).

### Experiment 2: Effects of circulating TNF-α on carotid sinus nerve afferent activity

Animals were anesthetized with isoflurane (Induction 5% and maintenance 2.5% in 100% O_2_) and subjected to femoral artery and vein catheterizations for arterial blood pressure (ABP) monitoring and drug administration, respectively, using polyethylene catheters (PE-50 attached to PE-10, Becton Dickinson, Sparks, MD, USA). Next, through a midline neck incision, the trachea was cannulated, and animals were artificially ventilated with a rodent ventilator (model 7025, Ugo Basile, Gemonio, VA, Italy). End-tidal CO_2_ was maintained between 4 - 5% (Capstar-100 carbon dioxide analyzer, CWE, Ardmore, PA, USA) by adjusting tidal volume (0.7 - 0.8 mL/100 g of body weight) and respiratory rate (60 - 80 bpm). Isoflurane was slowly replaced with urethane anesthesia (1.2 - 1.4 g/kg of body weight, IV) given over 20 - 25 minutes. Then, O_2_ concentration in ventilated air was switched to 50% O_2_ (balance N_2_) and this condition was kept until the end of the experiments. This slightly hyperoxic concentration was chosen because it ensures a stable preparation without silencing carotid body activity as 100% O_2_ would do (Kim et al., 2018; Schultz et al., 2007) and to avoid any period of hypoxia during the experimental protocol.

Then, animals were prepared for recordings of CSN afferent activity. The left carotid sinus nerve was identified, carefully isolated, and cut centrally at its junction to the glossopharyngeal nerve. CSN activity was recorded using bipolar suction electrodes and signals were filtered (100 - 3000 Hz), amplified (10,000 X) and digitally sampled (10 kHz). After baseline recordings, TNF-α (500 ng in 0.5 mL sterile saline, IV; PeproTech, Rocky Hill, NJ, USA) was administered and CSN activity was recorded for additional 2 hours. This dose was chosen based on previous works studying the effects of TNF-α on renal SNA in vivo (Zhang et al. 2003, Wei et al. 2015). Reliability of CSN activity was confirmed at the end of experiments by a robust increase in electrical activity during the exposure to hypoxia (10% O_2_) for 60 - 90 seconds.

### Experiment 3: Neuroanatomical identification of carotid body-related central sympathoexcitatory pathways activated by circulating TNF-α

First, the animals were anesthetized with a mixture of ketamine (80 mg kg^−1^, IP; União Química Farmacêutica Nacional S/A, Embu-Guaçu, SP, Brazil) and xylazine (8 mg kg^− 1^, IP; Hertape Calier Saúde animal S/A, Juatuba, MG, Brazil), and placed in a stereotaxic frame (David Kopf instruments, Tujunga, CA, USA). The retrograde tracer FluoroGold (FG, 2%, Fluorochrome, Denver, CO, USA) diluted in artificial cerebrospinal fluid (aCSF) was then bilaterally injected (40 nL) into the RVLM. Microinjections were performed with a pressure microinjector (Picospritzer III, Parker Hannifin, Hollis, NH, USA) using glass micropipettes. After each injection, the micropipette was kept in place for 2 minutes to prevent FG reflux. The coordinates used to target the RVLM were: 3.5 mm caudal from Lambda, 1.8 - 2.0 mm lateral from the midline, and 9.4 mm ventral from the skull surface. After injections, the skin incisions were sutured and the animals received anti-inflammatory ketoprofen (3 mg kg^−1^, SC) and antibiotics penicillin (50,000 IU, IM). This treatment was repeated every 24 hours for 3 days.

After 6 days recovery, animals were subjected to bilateral carotid body ablation (CB-X group) or Sham procedure (Sham group) and femoral artery/vein catheterizations. Carotid body ablation was performed by combining two previously described methods (Katayama et al., 2015; Pijacka et al., 2018). Briefly, animals were anesthetized with ketamine/xylazine as previously described. The carotid body arteries were ligated and cut, followed by surgical removal of the carotid bodies on both sides. In this procedure, the carotid sinus nerve is maintained intact, preserving carotid baroreflex function (Pijacka et al., 2018). Sham procedure consisted in isolation of carotid body arteries and carotid bodies, but these structures were kept intact. Neck incisions were closed with sutures. Femoral artery/vein catheters were tunneled subcutaneously, exteriorized and fixed in the interscapular region as previously described (Katayama et al., 2019). After surgeries, animals were treated with antibiotics and anti-inflammatory for 3 days as described before. To maintain catheters patency, arterial and venous catheters were flushed every day with heparinized saline (arterial: 500 U/mL, venous: 40 U/mL). Three days after surgery, ABP was recorded in unanesthetized rats under baseline conditions and in response to potassium cyanide (KCN; 40 ug/animal, IV) to verify the efficacy of carotid body ablation in CB-X group and the integrity of carotid bodies in SHAM group. Successful bilateral carotid body ablation was confirmed by the lack of cardiovascular responses to KCN (figure supplement 2A – B). Rats were allowed to recover for 3 days before the next experimental protocol.

On the day of the experiment (12 days after FG microinjections), rats were administered with TNF-α (500 ng in 0.5 mL sterile saline, IV) and left undisturbed for 2 hours. Next, rats were deeply anesthetized with urethane (IV) and transcardially perfused with PBS followed by PFA. Brains were collected and fixed in PFA for 12 hours at 4° C. Brains were then transferred to 20% sucrose solution and maintained at 4° C until the tissue sinks. Finally, brains were frozen in Tissue Freezing Medium (Triangle Biomedical Sciences, Durham, NC, USA) and coronal brain slices (30 µm) containing the cNTS and the RVLM were obtained on a cryostat. The RVLM sections were mounted on microscope slides (Superfrost Plus, Fisher Scientific, Pittsburgh, PA, USA) and used to confirm the location of FluoroGold microinjections within RVLM region (From 12.48 mm to 12.00 mm caudal to bregma, ventral to the compact formation of the Nucleus Ambiguus) accordingly to the rat brain in stereotaxic coordinates atlas (Paxinos and Charles Watson, 2007). The cNTS sections were stored in cryoprotectant solution at −20° C until processing for c-FOS and VGluT2 immunofluorescence as described below.

Briefly, sections were first rinsed in 0.1 M PBS for 10 minutes followed by incubation in blocking solution (0.1 M PBS, 10% normal horse serum, and 0.3% Triton X-100) for 20 min at room temperature. After rinsing 3 x 10 minutes in 0.1 M PBS at room temperature, slides were incubated in primary antibodies for c-FOS (1:1000, rabbit anti-c-FOS polyclonal antibody, product #sc-52, Santa Cruz Biotechnology, Santa Cruz, CA, USA) and for VGluT2 (1:2000, guinea pig anti-VGluT2 polyclonal antibody, product #AB2251-I, Millipore, Temecula, CA, USA) for 1 hour at room temperature plus 36 hours at 4° C. After rinsing in PBS, slides were incubated in secondary antibodies against rabbit (1:400, donkey anti-rabbit Alexa Fluor 594, product #A-21207, Molecular Probes-Life Technologies, Eugene, OR, USA) and against anti-guinea pig (1:400, donkey anti-guinea pig Alexa Fluor 488, product #706-545-148, Jackson ImmunoResearch Inc, West Grove, PA, USA) for 4 hours at room temperature. Slides were rinsed in PBS, the excess liquid was drained, mounting medium (Fluoromount, product # F4680, Sigma, St. Louis, MO, USA) was dropped on the tissue and slides were covered with glass coverslips (Fisherfinest, product #125485M, Fisher Scientific).

Images were acquired using a laser scanning confocal microscope (LSM800 with airyscan, Zeiss, Jena, TH, Germany). Quantification of retrograde labeled FG cells, c-FOS positive cells, FG/c-FOS cells, and FG/c-FOS/VGluT2 cells within the cNTS were performed on brainstem sections from three different rostro-caudal levels (between 14.76 mm to 14.04 mm caudal to bregma). These levels were chosen based on studies demonstrating NTS regions that are activated after carotid body stimulation (Cruz et al., 2010; Kline et al., 2010; Malheiros-Lima et al., 2020). As anatomical landmarks to identify the cNTS levels, we used: the area postrema, the central canal, the gracile nucleus, and the hypoglossal nucleus. Cell counting was performed on ImageJ software (U.S. National Institutes of Health, Bethesda, MD, USA) and representative figures were prepared using the Zen 2 software (Blue edition, Zeiss).

### Experiment 4: Sympathetic responses to circulating TNF-α in Sham and carotid body-ablated (CB-X) rats

Animals were anesthetized, subjected to femoral artery/vein catheterizations, tracheotomized and continuously ventilated with 50% O_2_ (balance N_2_) as described in *Experiment 2.* Next, animals were subjected to bilateral carotid body ablation or sham surgery as described in *Experiment 3*. All animals were then prepared for simultaneous recordings of lumbar, renal, and splanchnic sympathetic nerve activity (SNA). Lumbar sympathetic nerve was isolated through a midline laparotomy and retraction of vena cava, while renal and splanchnic sympathetic nerves were isolated through a retroperitoneal incision and careful retraction of the left kidney. Each sympathetic nerve was placed on a bipolar stainless-steel electrode and insulated with KWIK-SIL (World Precision Instruments, Sarasota, FL, USA). The raw SNA signals were filtered (100 - 1000 Hz), amplified (10,000 X) using biological amplifiers (P511 AC Amplifier, Grass Technologies, Warwick, RI, USA), displayed on oscilloscopes (TDS 2022, Tektronix, Beaverton, OR, USA) and digitally sampled (2 kHz) by a data acquisition system (Micro 1401, Cambridge Electronic Design Limited). All incisions were closed with surgical clips (Fine Science Tools, Foster City, CA, USA).

After stabilization (∼30 minutes after the end of surgical procedures), baseline recordings of ABP, lumbar, renal and splanchnic SNA were performed. Next, TNF-α (500 ng in 0.5 mL sterile saline, IV) was administered and ABP and SNA were recorded for additional 2 hours. At the end of the experiments, carotid body ablation was confirmed by the absence of blood pressure and SNA responses to KCN (40 ug/animal, IV) and these results are presented in figure supplement 3A – B.

### Experiment 5: Plasma and spleen levels of pro-inflammatory cytokines following intravenous TNF-α in SHAM, CB-X and SPL-X rats

Rats were anesthetized with ketamine/xylazine and prepared accordingly one of the following experimental groups: 1) CB-X: Animals were subjected to bilateral ablation of the carotid bodies; 2) SPL-X: Animals were subjected to splanchnic denervation through celiac ganglionectomy as previously reported in the literature (Asirvatham-Jeyaraj et al., 2021; Li et al., 2010). Briefly, after a midline laparotomy, the visceral organs were gently retracted, and the celiac ganglion was localized and surgically removed using blunt forceps. 3) SHAM: The carotid bodies and the celiac ganglion were localized and manipulated, but these structures were kept intact. All animals were allowed to recover for 6 days. Next, animals were subjected to femoral artery/vein catheterizations. The efficacy of carotid body ablation in CB-X group and the integrity of carotid bodies in SHAM and SPL-X groups was verified three days after catheterizations and these results are presented in figure supplement 4A – B. Then, rats were allowed to recover for additional 3 days before the experimental protocol.

The experimental protocol consisted in the administration of TNF-α (500 ng, IV) or vehicle (sterile saline, IV) in SHAM, CB-X and SPL-X rats. After 2 hours, rats were deeply anesthetized for tissue collection. Blood was collected into EDTA-containing tubes, centrifuged (1500 rpm for 10 min) at 4° C and the plasma was aliquoted and stored at −80° C. Spleen was harvested, flash frozen using liquid nitrogen and stored at −80° C.

The spleen samples were homogenized in PBS using a Polytron tissue homogenizer, and then centrifuged at 10,000 rpm for 2 min at 4 °C. The plasma and splenic levels of cytokines were quantified by enzyme-linked immunosorbent assay (ELISA) using commercial kits DuoSet ELISA Development Systems (R&D Systems, Minneapolis, MN, USA) for TNF-α (catalog #DY510), IL-6 (catalog #DY506), and IL-10 (catalog #DY522) and following the user manual instructions. The results were expressed as cytokine concentration in pg/mL and pg/mg of protein, based on standard curves, respectively. Spleen norepinephrine was measured as previously described (Garofalo et al., 1996) using HPLC (LC20AT-Shimadzu Proeminence) coupled to an electrochemical detector (Decade Lite-Antec Scientific) with a 5-μm Spherisorb ODS-2 reversed-phase column (Sigma-Aldrich) and the results were expressed as norepinephrine concentration in ng/g of tissue.

### Statistical analysis

All statistical analyses were performed using IBM SPSS Statistics (version 25, IBM corporation). Data are reported as means ± SEM. The significance level was set at p < 0.05, unless otherwise stated. No outliers were found as assessed by boxplot analyses. *Experiment 2*: To examine differences between means within the same group over time, the one-way repeated measures analysis of variance (ANOVA) followed by post hoc analysis with Bonferroni adjustment was performed. The normal distribution of the data was verified and confirmed by the Shapiro-Wilk test, and the Mauchly’s test indicated that the assumption of sphericity has not been violated. *Experiment 3*: To determine differences between two groups at a single time-point, we first assessed the distribution of the data using the Shapiro-Wilk test and the homogeneity of variances using the Levenés test. For normally distributed data with homogenous variances, an unpaired two-tailed Student’s *t*-test was performed. In cases in which the data was normally distributed but the assumption of homogeneity of variances was violated, an unpaired two-tailed Welch’s *t*-test was used. When data was not normally distributed, an unpaired two-tailed Mann-Whitney *U*-test was applied. *Experiment 4*: To determine group x time interactions, a two-way repeated measures ANOVA was conducted. In this case, the normal distribution of the studentized residuals was verified and confirmed by the Shapiro-Wilk test. The sphericity for the interaction term was assessed by the Mauchly’s test. When the assumption of sphericity was violated, the Greenhouse-Geisser correction was used and the estimated epsilon (ε) value was reported. Once statistically significant group x time interactions were detected, simple main effects of group were analyzed using repeated measures general linear models with Bonferroni adjustment. *Experiment 5*: To examine group x treatment interactions and main effects of group, a two-way ANOVA with Bonferroni post hoc was performed. The distribution of the residuals was examined by the Shapiro-Wilk test. The homoscedasticity was analyzed by the Levenés test and by plotting the residuals against the predicted values in a simple scatterplot. If the assumptions of normal distribution and/or homoscedasticity were violated, the dependent variable was log-transformed when appropriate. When both the assumptions of normality and homoscedasticity (requirements for two-way ANOVA) were not satisfied even after transformation, a Kruskal-Wallis followed by Mann-Whitney *U*-tests for pairwise comparisons between groups were performed. In these cases, a Bonferroni adjustment to alpha values was applied and the statistical significance was accepted at the p < 0.003 level. In cases in which only the assumption of homoscedasticity was violated, a Welch’s ANOVA followed by a Games-Howell post hoc was used to compare groups.

## Acknowledgments

The authors thank Lilian do Carmo Heck for the excellent technical assistance. This work was funded by the São Paulo Research Foundation (FAPESP; grants #2019/11196-0 and #2015/23467-7), CNPq, and PROPE-UNESP.

## Author contributions

P.L.K. and E.C. conceived and designed research. P.L.K. performed all in vivo experiments. P.L.K. and I.P.L. performed immunofluorescence and in situ hybridization. A.K. and J.P.M.L. performed ELISA. L.C.C.N. performed HPLC measurements. P.L.K., I.P.L., A.K., and J.P.M.L. analyzed data. P.L.K., I.P.L., A.K., D.B.Z, and E.C. interpreted data. P.L.K. and A.K. drafted the manuscript. P.L.K., I.P.L., A.K., J.P.M.L., F.Q.C., L.C.C.N., J.V.M., D.B.Z., D.S.A.C., and E.C. edited and revised the manuscript. P.L.K., I.P.L., A.K., J.P.M.L., F.Q.C., L.C.C.N, J.V.M., D.B.Z., D.S.A.C., and E.C. read and approved the final version of the manuscript.

## Materials & correspondence

Correspondence and requests for materials should be addressed to P.L.K. and/or E.C.

## Supplementary Information

**Figure supplement 1.**
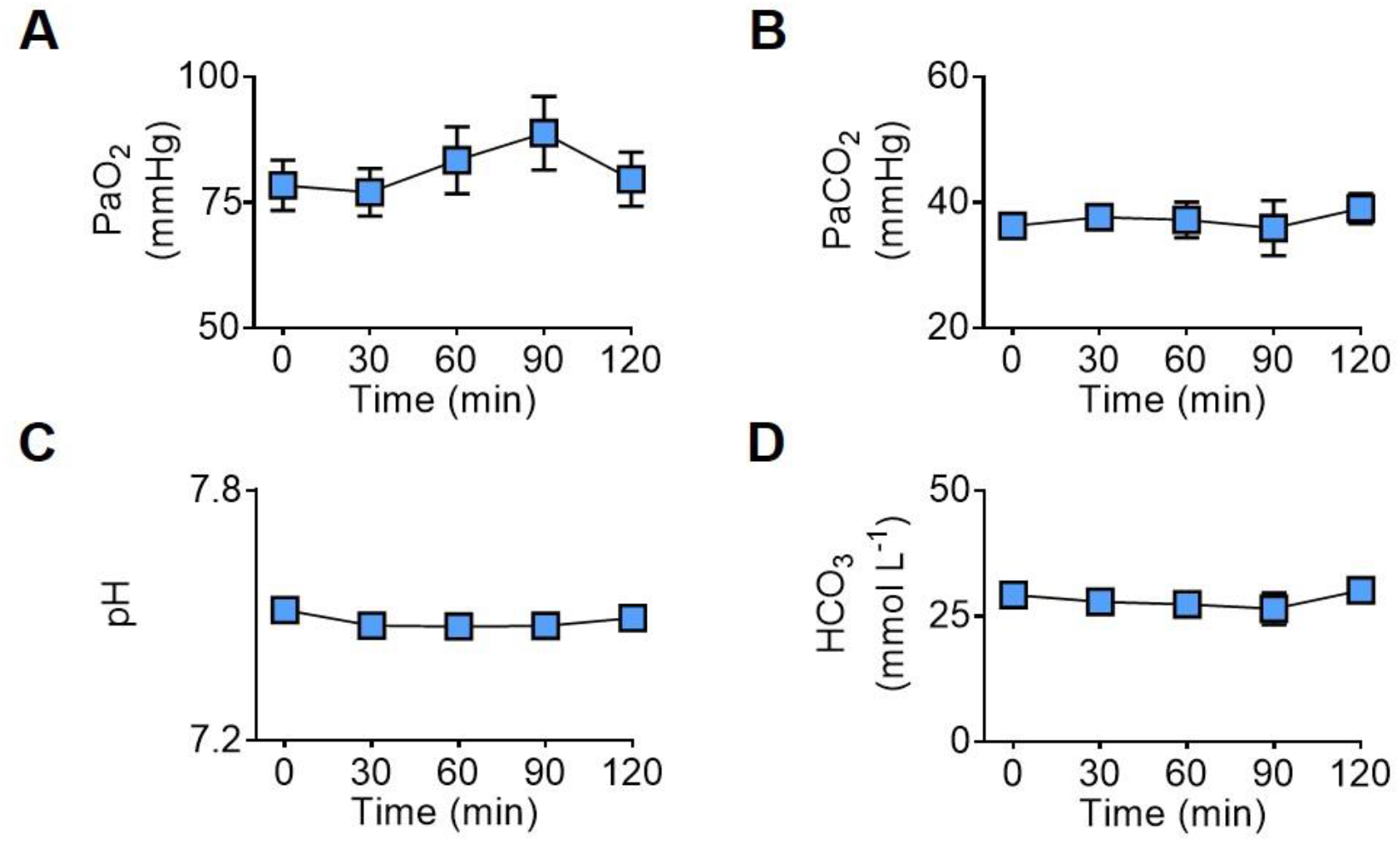
Acute intravenous TNF-α does not affect arterial blood gases, pH, and bicarbonate. All measures were performed using a I-STAT device with CG4+ cartridges (Abbott, Abbott Park, IL, USA). **A – D.** Summary data (n = 5) showing that the intravenous administration of TNF-α (500 ng) did not affect the partial pressure of oxygen (PaO_2_), the partial pressure of carbon dioxide (PaCO_2_), the pH, and the bicarbonate (HCO_3_^−^) concentration in the arterial blood of unanesthetized, spontaneously breathing rats. One-way repeated measures ANOVA: PaO_2_, *F*_(1.525, 6.102)_ = 1.659, p = 0.259, ε = 0.381; PaCO_2_, *F*_(4,16)_ = 0.370, p = 0.826; pH, *F*_(4,16)_ = 2.838, p = 0.059; HCO_3_^−^, *F*_(1.879, 7.515)_ = 1.207, p = 0.347, ε = 0.470. Data are means ± SEM.

**Figure supplement 2.**
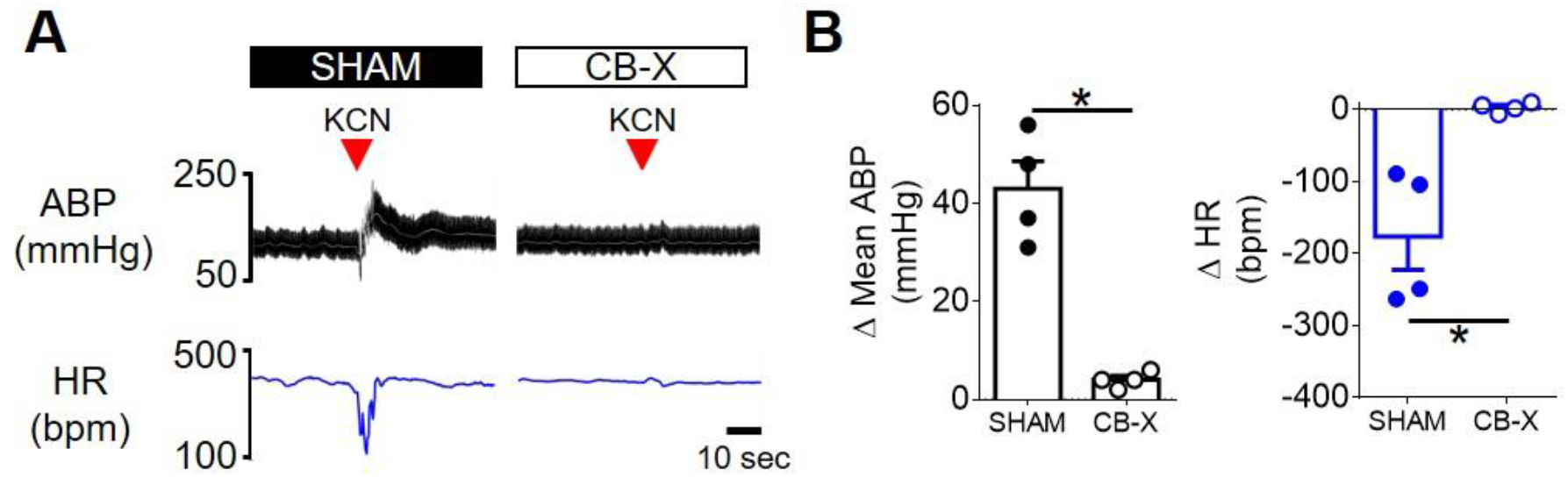
Verification of carotid body ablation in *experiment 3*. **A.** Representative tracings of arterial blood pressure (pulsatile ABP, black; mean ABP, white) and heart rate (HR; blue) of a SHAM rat (left) and of a CB-X rat (right) in response to KCN (red arrowhead, 40 µg, IV) under unanesthetized conditions. **B.** Summary data showing the peak changes in mean ABP and HR in response to KCN from SHAM (filled symbols, n = 4) and CB-X (open symbols, n = 4) rats. The cardiovascular responses to carotid body stimulation by intravenous KCN were abolished in CB-X rats, confirming the efficacy of bilateral carotid body ablation: Δ mean ABP (SHAM, 43 ± 6 mmHg; CB-X, 4 ± 1 mmHg; *t*(3.128) = 6.912, p = 0.005, Welch’s *t*-test), Δ HR (SHAM, −176 ± 46 bpm; CB-X, 3 ± 3 bpm; *t*(3.033) = −3.866, p = 0.03, Welch’s *t*-test). *p < 0.05. Data are means ± SEM.

**Figure supplement 3.**
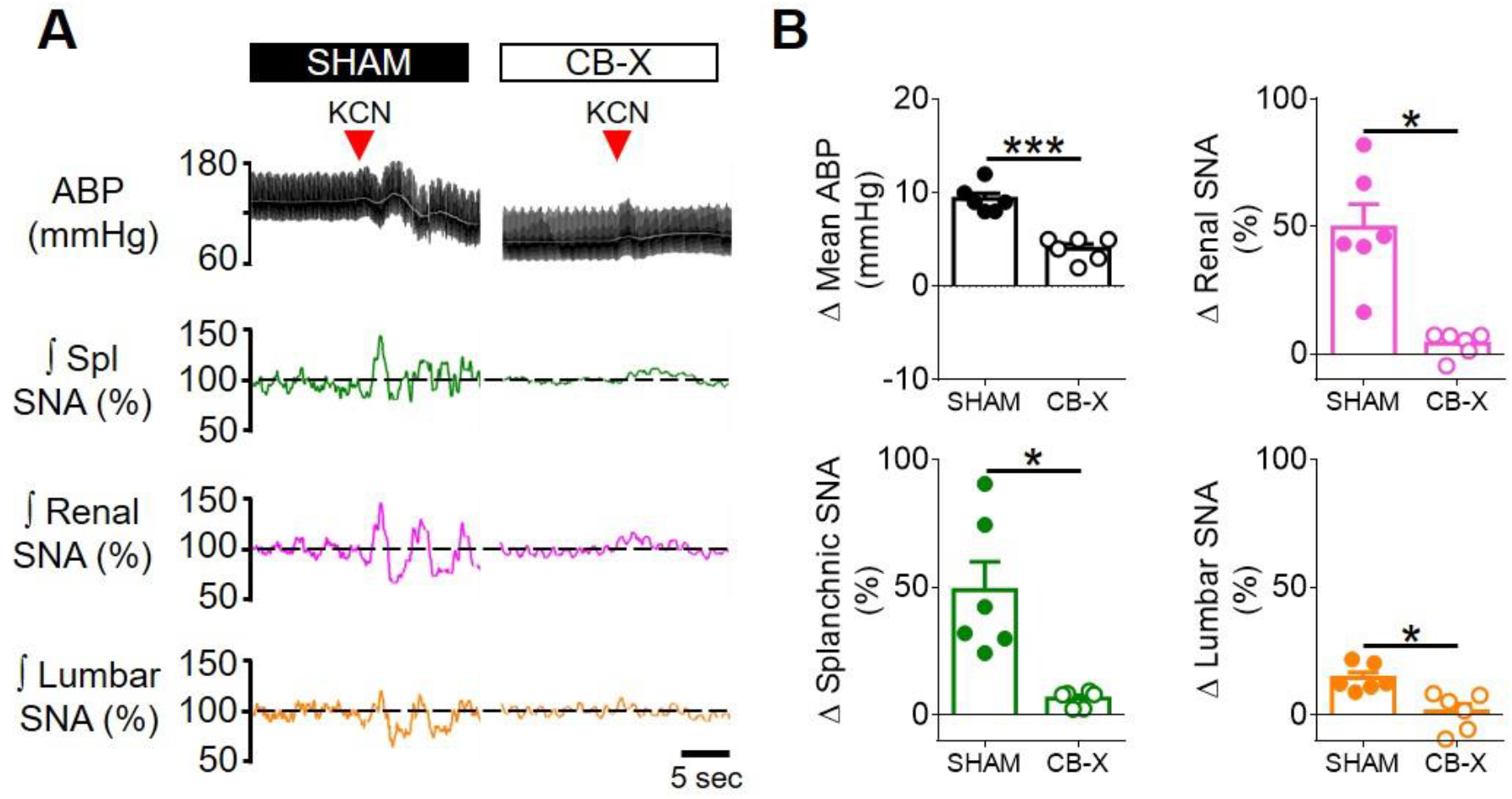
Verification of carotid body ablation at the end of *experiment 4*. **A.** Representative tracings of arterial blood pressure (pulsatile ABP, black; mean ABP, white), splanchnic SNA (Spl; green), renal SNA (magenta) and lumbar SNA (orange) of a SHAM rat (left) and of a CB-X rat (right) in response to KCN (red arrowhead, 40 µg, IV) under anesthetized conditions. **B.** Summary data showing the changes in mean ABP, Splanchnic SNA, Renal SNA and Lumbar SNA in response to KCN from SHAM (filled symbols, n = 6) and CB-X (open symbols, n = 6) rats. For each rat, baseline rectified and integrated SNA was normalized to 100% and the peak changes in response to KCN were calculated. The sympathetic and blood pressure responses to KCN were significantly attenuated in CB-X rats, confirming the efficacy of bilateral carotid body ablation: Δ Spl SNA (SHAM, 48 ± 11 %; CB-X, 7 ± 1 %; *U* = 0, _Z_ = −2.882, p = 0.002, Mann-Whitney *U*-test), Δ Renal SNA (SHAM, 50 ± 9 %; CB-X, 4 ± 2 %; *t*(5.452) = 4.815, p = 0.004, Welch’s *t*-test), Δ Lumbar SNA (SHAM, 15 ± 2 %; CB-X, 1 ± 3 %; *t*(10) = 3.547, p = 0.005, Student’s *t*-test), and Δ Mean ABP (SHAM, 9 ± 1 mmHg; CB-X, 4 ± 1 % ; *t*(10) = 6.644, p < 0.001, Student’s *t*-test). *p < 0.05 and ***p < 0.001. Data are means ± SEM.

**Figure supplement 4.**
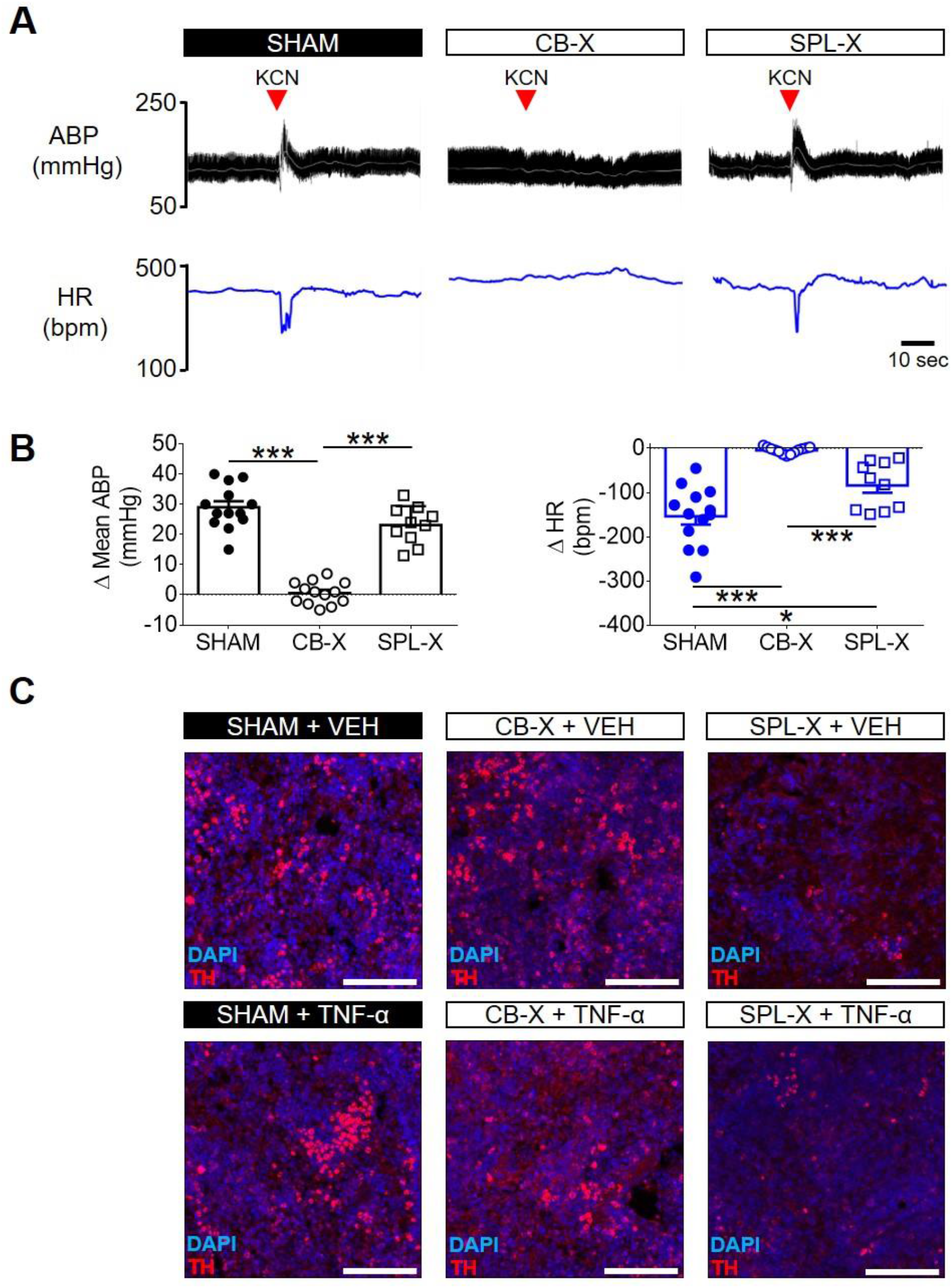
Verification of carotid body ablation and splanchnic sympathetic denervation in *experiment 5*. **A.** Representative tracings of arterial blood pressure (pulsatile ABP, black; mean ABP, white) and heart rate (HR; blue) of a SHAM rat (left), of a CB-X rat (middle), and of a SPL-X rat (right) in response to KCN (red arrowhead, 40 µg, IV) under unanesthetized conditions. **B.** Summary data showing the peak changes in mean ABP (black graphs, left) and HR (blue graphs, right) in response to KCN from SHAM (filled circles, n = 13), CB-X (open circles, n = 13), and SPL-X (open squares, n = 10) rats. The cardiovascular (mean ABP and HR) responses to carotid body stimulation by intravenous KCN were abolished in CB-X rats, confirming the efficacy of bilateral carotid body ablation: Δ mean ABP (SHAM, 29 ± 2 mmHg; CB-X, 1 ± 1 mmHg; SPL-X, 23 ± 2) and Δ mean HR (SHAM, −154 ± 19 bpm; CB-X, −4 ± 2 bpm; SPL-X, −84 ± 17). Regarding Δ mean ABP, a one-way ANOVA detected statistically significant differences between groups, *F*_(2, 33)_ = 83.134, p < 0.001. Subsequent post hoc analysis with a Bonferroni adjustment revealed that the mean difference in Δ mean ABP between CB-X and SHAM rats was statistically significant (−28 mmHg, 95% CI [−34, −23], p < 0.001) as well as the mean difference in Δ mean ABP between CB-X and SPL-X rats (−22 mmHg, 95% CI [−29, −16], p < 0.001). Regarding Δ HR, a Welch ANOVA detected statistically significant differences between groups, *F*_(2, 14.078)_ = 40.040, p < 0.001. Games-Howell post hoc analysis revealed that the mean difference in Δ HR between CB-X and SHAM rats was statistically significant (149 bpm, 95% CI [99, 200], p < 0.001) as well as the mean difference in Δ mean HR between CB-X and SPL-X rats (79 bpm, 95% CI [33, 126], p = 0.003). In addition, the mean difference in Δ mean HR between SHAM and SPL-X rats was also statistically significant (−70 bpm, 95% CI [−133, −7], p = 0.029). *p < 0.05 and ***p < 0.001. Data are means ± SEM. **C.** Representative images of spleen sections from one animal of each group obtained at the end of *experiment 5* and processed for nuclear staining (DAPI, blue) and tyrosine hydroxylase (TH, red). Note that TH staining is substantially less pronounced in the animals subjected to splanchnic sympathetic denervation (SPL-X + VEH, upper right panel; and SPL-X + TNF-α, bottom right panel) as compared to SHAM (SHAM + VEH, upper left panel; and SHAM + TNF-α, bottom left panel) and CB-X (CB-X +VEH, upper middle panel; and CB-X + TNF-α, bottom middle panel). VEH, vehicle. Scale bars: 100 µm.

